# Dystrophin modulates focal adhesion tension and YAP-mediated mechanotransduction

**DOI:** 10.1101/2021.08.05.455167

**Authors:** Maria Paz Ramirez, Michael J.M. Anderson, Lauren J. Sundby, Anthony R. Hagerty, Sophia J. Wenthe, James M. Ervasti, Wendy R. Gordon

## Abstract

Dystrophin is an essential muscle protein that contributes to cell membrane stability by linking the actin cytoskeleton to the extracellular matrix. The absence or impaired function of dystrophin causes muscular dystrophy. Focal adhesions are mechanosensitive adhesion complexes that also connect the cytoskeleton to the extracellular matrix. However, the interplay between dystrophin and focal adhesion force transmission has not been investigated. Using a bioluminescent tension sensor, we measured focal adhesion tension in transgenic C2C12 myoblasts expressing wild type (WT) dystrophin, a non-pathogenic SNP (I232M), or two missense mutations associated with Duchenne (L54R), or Becker muscular dystrophy (L172H). We found that myoblasts expressing WT or nonpathogenic I232M dystrophin showed increased focal adhesion tension compared to non-transgenic myoblasts, while myoblasts expressing L54R or L172H dystrophin presented with decreased focal adhesion tension. Moreover, myoblasts expressing L54R or L172H dystrophin showed decreased YAP activation and exhibited slower and less directional migration compared to cells expressing WT or I232M dystrophin. Our results suggest that disease-causing missense mutations in dystrophin may disrupt a cellular tension sensing pathway in dystrophic skeletal muscle.

## Introduction

Skeletal muscle is an organ under constant mechanical stress, even at rest. Dystrophin, a protein located beneath the muscle cell plasma membrane (sarcolemma)(Zubrzycka-Gaarn et al. 1988), is a key cytoskeletal protein that contributes to maintaining muscle integrity(Hoffman, Brown, and Kunkel 1987; Rybakova, Patel, and Ervasti 2000; James M. Ervasti 2007). Dystrophin maintains the structural stability of the sarcolemma by linking the actin cytoskeleton to the extracellular matrix (ECM), via a transmembrane protein adhesion complex known as the dystrophin-glycoprotein complex (DGC)(J. M. Ervasti et al. 1990; J. M. Ervasti and Campbell 1991; Ohlendieck et al. 1991; Williams and Bloch 1999). The DGC is enriched at the costameres in striated skeletal muscle, which are subsarcolemmal protein assemblies that couple the force-generating sarcomeres to the sarcolemma (Pardo, Siliciano, and Craig 1983; Straub et al. 1992; Rybakova, Patel, and Ervasti 2000; James M. Ervasti 2003). Mutations in genes encoding different components of the DGC are the cause of various forms of muscular dystrophy, such as Duchenne and Becker muscular dystrophy (DMD and BMD, respectively). DMD is caused by the absence of dystrophin (Hoffman, Brown, and Kunkel 1987), while BMD is caused by decreased expression of a truncated and less functional dystrophin (Beggs et al. 1991; Hamed et al. 2005). Partial to total loss of dystrophin leads to sarcolemmal fragility (Menke and Jockusch 1991; Mokri and Engel 1998) and myofiber death (Tidball et al. 1995), which is outwardly manifested as muscle weakness (Goldstein and McNally 2010).

Another cytolinker complex found in skeletal muscle are integrin-based focal adhesions (FA), which connect the ECM to the cytoskeleton via talin and vinculin (Burridge and Chrzanowska-Wodnicka 1996; Wozniak et al. 2004). Cells sense, generate, and respond to forces through mechanosensitive protein complexes, such as those in FA. Mechanical signals are accumulated and transmitted via the cytoskeleton to mechanotransducer proteins to ultimately modulate gene expression that regulates cell behavior, in a process known as mechanotransduction (Chen 2008; Jaalouk and Lammerding 2009). Yes-associated protein 1 (YAP) is a central mechanotransducer under the control of FA activity (Lachowski et al. 2018; Dasgupta and McCollum 2019). Dysregulation of mechanical signals, force transmission between ECM and cytoskeleton and mechanotransducer activities have been shown to promote pathogenic phenotypes that promote muscular dystrophy (Kumar et al. 2004; Graham, Gallagher, and Cardozo 2015; Boppart and Mahmassani 2019). FA and the DGC interact via the actin cytoskeleton, yet the interplay between dystrophin and force transmission via FA has not been investigated. Furthermore, the effect of disease-causing dystrophin missense mutations in FA tension and mechanotransducer regulation has not been established.

Previous studies have shown that dystrophin contributes to the mechanical stability of muscle, and that loss of dystrophin leads to decreased tissue and cellular stiffness. For example, the sarcolemmal cytoskeleton of myofibers from dystrophin-null *mdx* mice are more compliant than healthy controls (García-Pelagio et al. 2011; Pasternak, Wong, and Elson 1995; Kumar et al. 2004). Muscle compliance is also increased in *mdx* mice compared to controls (Lopez et al. 2021). In addition, there is increasing data that YAP activity is aberrant in muscular dystrophy (Hulmi et al. 2013; Morikawa et al. 2017; Vita et al. 2018; Iyer et al. 2019). We hypothesize that dystrophin enhances links between the ECM and cytoskeleton in muscle cells, thus influencing FA tension and in turn, force transmission and regulation of mechanotransducer proteins, like YAP. Dystrophin mutations, then, likely lead to abnormal FA tension and mechanotransduction.

Here, we used molecular tension sensors to study the impact of dystrophin on FA tension sensing in C2C12 myoblasts transgenically expressing wildtype or mutant dystrophin. We show that stable expression of wildtype dystrophin increases FA tension compared to dystrophin-less non-transgenic control cells. However, DMD- or BMD-associated missense mutations act to decrease focal adhesion tensions.

Our results showed that decreased FA tension in the dystrophin-mutant cell lines also positively correlated to reduced YAP mechanosignaling and impaired cell migration, linking specific defects in mechanotransduction at the molecular level to muscle-level phenotypes that promote disease progression. Our novel data are the first to demonstrate a role for dystrophin FA tension and YAP-mediated mechanotransduction.

## Methods

### Cell Culture

C2C12 (ATCC; CRL-1772) and transgenic C2C12 cell lines were cultured and grown at 37 °C, 5% CO_2_ in Gibco Dulbecco’s Modified Eagle Medium (DMEM, Gibco) supplemented with 10% fetal bovine serum (FBS, ATCC), 1% penicillin/streptomycin mixture (HyClone; 15290), and 0.1% fungizone (Gibco). Cells were cultured on either plastic cell dishes or on Matek glass bottom plates previously coated with 12.5 ng/mL fibronectin solution in phosphate buffered saline (PBS 1X, Corning).

### BRET Imaging

Cells cultured on fibronectin coated glass bottom plates were washed with PBS and the DMEM culture media was replaced with warm FluoroBrite™ media (Gibco) supplemented with 10% FBS and 1:1000 Nano-Glo^®^ Luciferase Assay Substrate (Furimazine, Promega). First, the donor and acceptor emissions were imaged at 100x using an IX83 inverted microscope (Olympus) equipped with an iXon Ultra 888 EM-CCD camera (Andor), using a filter wheel with Semrock light filters FF01-460/60 (NanoLuc, donor) and Semrock FF01-536/40 (mNeonGreen, acceptor). Integration time was 45 s for both acquisitions. Then cells were imaged for mNeonGreen using a LED lighting source and a standard FITC cube, to obtain a regular fluorescent image of the FA.

### Immunostaining

C2C12 cells were cultured on fibronectin coated glass bottom plates. After 24 hr incubation, cells were fixed with 4% paraformaldehyde (Thermo Fisher Scientific) for 13 min at room temperature (RT) and immediately rinsed with RT PBS twice. Fixed cells were permeabilized and blocked with a PBS + 2% Bovine serum albumin (BSA, Fisher BioReagents) + 0.05% Triton X-100 (Fisher Scientific) blocking solution for 10 min at RT. Blocked cells were labeled with YAP1 (1:100, Abnova; 2F12) and paxillin (1:100, Abcam; Y113) antibodies in blocking buffer for 1 hr at RT, then washed three times with blocking buffer. The cells were then incubated for 45 min with secondary antibody diluted in blocking buffer (1:500, Alexa Fluor 568, Molecular Probes) and washed twice with blocking buffer and once with PBS. Finally, labeled cells were mounted with ProLong™ Diamond antifade with DAPI (Thermo Fisher Scientific).

Immunostained cells were imaged on an IX83 inverted microscope (Olympus) equipped with an iXon Ultra 888 EM-CCD camera (Andor), using a LED lighting source and standard FITC, TRITC and DAPI filter cubes. Image overlays were created using ImageJ (version 1.53h).

### Image Analysis

Acquired emission images were analyzed on ImageJ (version 1.53h). Stacked images were aligned using the Linear Stack with Alignment with SIFT plugin and then analyzed using the BRET-Analyzer plugin(Chastagnier et al. 2017). mNeonGreen images were used to generate FA masks using a machine learning tool, Ilastik (version 1.3.2). Ratiometric images generated by the BRET-Analyzer were masked and BRET distributions were obtained and visualized as 16 color LUT. To analyze FA tension gradients from the BRET ratiometric images, line scans were done across individual FA by manually using line tool, from the edge of the cell towards the centroid. Obtained intensity profiles were normalized to FA length. Cell and FA morphology analysis were done using the Analyze Particles Plugin.

Immunostained images were masked for the complete cell and for the nucleus; masks were generated using Ilastik. Using ImageJ, YAP cellular and nuclear intensity was measured, and then nuclear intensity was subtracted from cellular intensity to calculate cytoplasmic YAP intensity. From this data, nuclear to cytoplasmic YAP ratios were calculated.

### Western Blot Analysis

Harvested cells were weighted and lysed with a volume of lysis buffer proportional to cell pellet mass. Lysis buffer used was radioimmunoprecipitation assay buffer (RIPA, 10 mM Tris·Cl (pH 8.0), 1 mM EDTA, 0.5 mM EGTA, 1% Triton X-100, 0.1% sodium deoxycholate, 0.1% SDS, 140 mM NaCl) with added protease inhibitors (100 nM Aprotinin, 10 μM E-64, 10 μM Leupeptin, 1 mM Pepstatin A, 1 mM PMSF). Lysates sampled with phospho-specific antibodies were also lysed with phosphatase inhibitors (phosSTOP™, Roche). Lysates were then centrifuged at 20,000 g, and equal volumes of collected supernatant were separated on sodium dodecyl sulphate (SDS) polyacrylamide gel at 100 V for 30 min and 150 V for 1 h and transferred to a PVDF membrane at 0.8A for 1 h. Membranes were blocked for 1 h in either 5% milk in PBS 0.1% Tween (PBST) or 5% BSA in Tris-buffered saline (TBS) 0.1% Tween (TBST), depending on the primary antibody used. Blocked membranes were incubated with primary antibodies overnight at 4°C. Primary antibodies diluted with milk PBST were GFP (1:1,000 Cell Signaling; 2956), pan actin C4 (1:5,000 Seven Hills Bioreagents; LMAB-C4), α-tubulin (1:1000, Sigma-Aldrich; DM1A), utrophin (1:50, DHSB; Mancho3) and YAP (1:500, Abnova; 2F12). Primary antibodies diluted with BSA TBST were vinculin (1:1000, Sigma-Aldrich; V9131), paxillin (1:100, Abcam; Y113), α-dystroglycan (1:500, Millipore; 05-298), β-dystroglycan (1:1,000, Thermo Scientific; pA5-34908) and P-YAP S127 (1:2000, Abcam; EP1675Y). Membranes were washed with PBST or TBST and then incubated with secondary antibody diluted in the appropriate blocking buffer. Secondary antibodies used were anti-mouse or anti-rabbit IgG Dylight 800 or IgG Dylight 700 (1:10,000, Cell Signaling). Secondary antibody signal was visualized on LI-COR’s Odyssey CLx Imaging System and band density was calculated with Image Studio™ Software.

### RT-qPCR

Cells were collected and their RNA was extracted using Aurum Total RNA Mini Kit (Bio-Rad; 732-6820). Total RNA concentration and purity was determined using a nanodrop spectrophotometer (Thermo Scientific). A standardized amount of RNA was transcribed into cDNA using the iScript Advanced cDNA Synthesis Kit (Bio-Rad; 170-8843), and cDNA was then amplified using the primers: GFP forward (TCCGCCATGCCCGAAGGCTA) and reverse (CCGTTCACCAGGGTGTCGCC), to detect transgenic dystrophin (GFP-dystrophin). Measurements were done relative to Hprt using the primers: forward (CCCTGGTTAAGCAGTACAGCCCC) and reverse (GGCCTGTATCCAACACTTCGAGAGG). cDNA was amplified using SsoAdvanced Universal SYBR Green Supermix (Bio-Rad; 172-5270) on the C1000 Touch Thermal Cycler (Bio-Rad) and then analyzed with the CFX Manager software (Bio-Rad).

### Wound Healing Assay

Confluent monolayers of C2C12 cells plated on fibronectin were gently washed with PBS and then scratched with a 200 μL micropipette tip to create a wound (cell free area) all along the center of the well. Then, cells were washed once with PBS and warm FluoroBrite™ media supplemented with 10% FBS was added. Scratched cells were incubated at 37 C and 5%CO2 and imaged every hour over 9 hours imaged on a bright field for 9 h every hour at 37 °C and 5% CO2 on an IX83 inverted microscope (Olympus) equipped with an iXon Ultra 888 EM-CCD camera (Andor) in a top stage incubator (STX, Tokai Hit). The percentage of wound closure was calculated as:

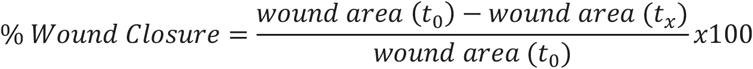

Then, the % wound closure over time was plotted and fitted the linear portion of the data to a y = mx linear fit, where x is the slope of the fitted line.

Inferred edge “speed” of wound closure was calculated as:

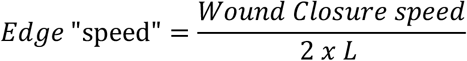

As the migration of the sheet of cells can effectively be considered one directional, only towards the center of the wound, the vertical velocity was assumed to be zero. Therefore, the area of the wound only changes by its width and not length (L), which is twice as slow as the change in wound size as both cell sheets contribute to wound closure.

The wound closure speed (μm2/time) was calculated by multiplying the fitted slope from the % wound closure over time with the initial wound size, as

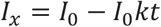

Where “I” is the wound size, k is the slope obtained from the %wound closure fit and t is time.

### Live Cell Imaging

5×10^3^ C2C12 cells were cultured on a Nunc™ glass base dish (Thermo Scientific) coated with fibronectin and cultured overnight. Cells were then washed with PBS and media was replaced with warm FluoroBrite™ supplemented with FBS that was incubated overnight at 37 °C, 5% CO_2_. Cells were imaged on a Delta Vision personalDV (GE Technologies), using a 10x/NA0.25 objective, in a closed environmental chamber at 37°C, with phase contrast illumination, for 4 h at 10 min intervals. Migration was tracked using the ImageJ Manual Tracking plugin and the xy track data was analyzed using the DiPer excel plugin as described by (Gorelik and Gautreau 2014). Cells that contacted other cells, came in and out of frame or that visibly divided during imaging were excluded for analysis.

### Proliferation Assay

8×10^2^ C2C12 cells were plated on 96 wells in triplicate for each measurement day (4 days) and cultured overnight with DMEM. The next day, CellTiter-Glo™ substrate, media and cell plate were left to equilibrate at room temperature. Cells were lysed with a 1:1 solution of substrate and DMEM for 10 min. The cell and substrate mixture was then transferred to an opaque white 96 well plate and luminescence was measured using the LMax II 384 Luminometer (Molecular Devices). A blank of DMEM was included, and was subtracted from the values acquired for each measurement day.

### Statistical Analysis

All statistical calculations were performed using the Graphpad Prism 8 software. All data are presented as mean ± SEM. Plots showing layered data were created as superplots described by (Lord et al. 2020). Unpaired two-tailed t-test was performed to compare the means of variables between VinTL and VinTS cells (same cell line), performed at α = 0.05. Unpaired one-way ANOVA analysis was performed to compare the means of three or more populations, with α = 0.05 to determine significance. Significance was confirmed with a Tukey’s post hoc test, performed at α = 0.05. Unpaired two-way ANOVA analysis was performed to determine the significant differences between populations to a dependent variable (e.g. time), at α = 0.05.

## Results

### Increased focal adhesion tension in myoblasts stably expressing dystrophin

In order to measure tensions in muscle cells harboring variable dystrophin levels and mutations, we use previously characterized transgenic C2C12 myoblasts lines(Talsness, Belanto, and Ervasti 2015) that stably express wildtype dystrophin, or dystrophins with missense mutations associated with DMD and BMD(Talsness, Belanto, and Ervasti 2015; Prior et al. 1993; Hamed et al. 2005). These transgenic cell lines overcome the limitation of using C2C12 cell lines, which do not exhibit detectable levels of dystrophin until they are differentiated and cultured for long periods. In order to probe ECM to cytoskeleton tensions at a molecular level using focal adhesions as a proxy, we use an established bioluminescence-based molecular tension sensor-BRET-TS-genetically-encoded in the focal adhesion protein vinculin (Grashoff et al. 2010; LaCroix et al. 2018; Aird et al. 2020). BRET-TS consists of a flexible polypeptide “spring” derived from the spider silk flagelliform domain (GPGGA)_8_ flanked by NanoLuc donor and mNeonGreen acceptor proteins(Fig. 1A)(Aird et al. 2020). BRET-TS is localized between the head domain of vinculin − V_h_ − that binds to talin and the tail domain of vinculin − V_t_ − that binds to actin (VinTS, Fig. 1B). When vinculin is under high tension, BRET ratios decrease as the flexible module stretches, separating the donor and acceptor BRET pair (Fig. 1B). A truncated version of the tension sensor that cannot experience tension because it lacks the actin binding tail (VinTL, Fig. 1B) was used as a control.(Grashoff et al. 2010).

**Figure 1.**
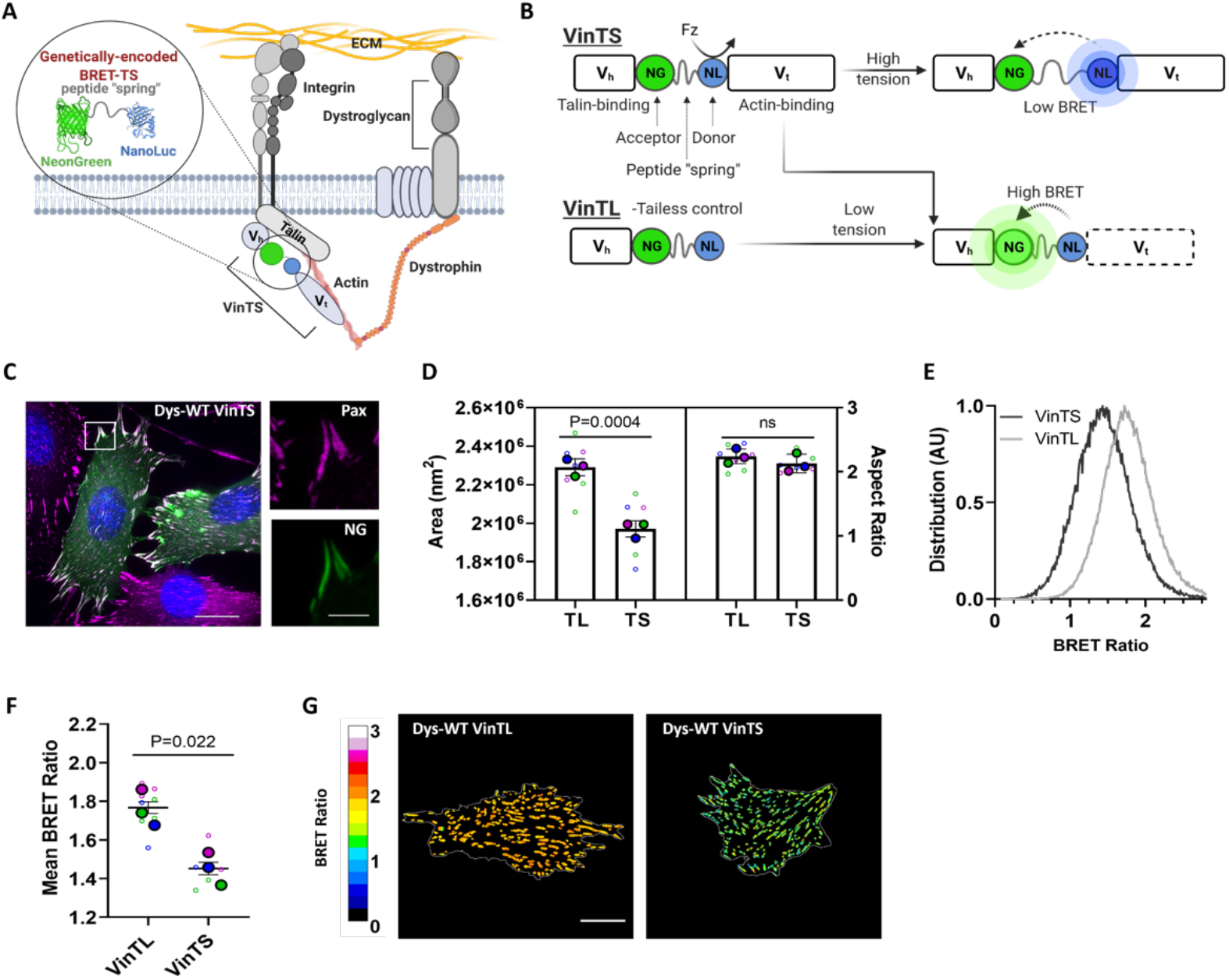
Vinculin BRET Tension Sensor in Focal Adhesions. **A.** Schematic of Bioluminescent Resonance Energy Transfer Tension Sensor (BRET-TS) inserted into vinculin in focal adhesions (VinTS) and **B.** of the BRET efficiencies of VinTS and VinTL under in the presence and absence of forces. BRET-TS is inserted between the talin-binding head domain (Vh) and the actin binding tail domain (Vt). The DGC is depicted to the right of a focal adhesion, linked together via actin. Other DGC components are left nameless for simplicity. The addition of furimazine (Fz) excites the energy donor NanoLuc (NL). Efficiency of energy transfer to the acceptor NeonGreen (NG) is dependent on the load experienced by the tension sensor. **C.** Representative image of C2C12 myoblasts transfected with VinTS. Fixed cells expressing VinTS (NG) were immunostained with paxillin (magenta) and DAPI (blue) to indicate focal adhesions and nucleus. **D.** Focal adhesion morphology comparison (N=3) of myoblasts transfected with VinTL and VinTS. Large dots denote the mean of independent experiments and smaller dots denote the mean of individual focal adhesion measurements per cell. Each experiment and corresponding individual measurements are colored the same. **E.** Distribution and **f.** Distribution mean of app BRET efficiency (N=3) for VinTS and VinTL in transfected myoblasts. **G.** Representative images of apparent BRET efficiency of VinTS and VinTL in myoblasts. All data shown is from Dys-WT myoblasts. Scale bars indicate 20 μm and 5 μm for zoom. Data analyzed via an unpaired 2-tailed t-test; ns, not significant. All error bars represent SEM.

We first aimed to validate the activity of the tension sensor in C2C12 myoblasts and ensure that it did not disrupt focal adhesion formation. Our overall workflow involves transient transfection of vinculin tension sensors into the transgenic cell lines and live-cell imaging to measure the extension of a flexible peptide via the BRET ratio of mNeonGreen to NanoLuc emission. VinTS correctly localized to FA in transiently transfected myoblasts (Fig. 1C and Supp. 1). Vinculin levels remained constant across myoblasts expressing dystrophin and non-transgenic myoblasts (henceforth referred as NTg) (Supp. 2), with no significant difference in expression of the tensor sensor (Supp. 3). These results confirm that the tension sensor does not alter vinculin localization or protein levels. VinTL expressing myoblasts formed larger FA than VinTS (Fig. 1D and Supp. 4), consistent with observations made by others (Grashoff et al. 2010; Humphries et al. 2007). The shape (aspect ratio) of VinTS and VinTL FA and overall cell morphologies were not different (Fig. 1D, Supp. 4, Supp 5).Therefore, tension sensor, vinculin levels, FA shape and cell morphology were all constant between VinTL and VinTS.

To assess whether VinTS sensed tension at FA, we performed ratiometric BRET imaging on VinTS and VinTL encoded myoblasts expressing wildtype dystrophin (WT-Dys). As expected, we found that cells transfected with VinTS showed significantly lower BRET ratios when compared to VinTL transfected cells (Fig. 1E-G), confirming that VinTS is under greater mechanical tension than VinTL. Interestingly, we did not observe tension gradients along FA as we and others have previously reported (Supp. 6-7)(Sarangi et al. 2017; Rothenberg et al. 2018; Aird et al. 2020). Altogether, these results validate the use of VinTS as a tension sensor to study levels of tension transmitted across vinculin in FA of C2C12 myoblasts.

When compared to NTg cells, which do not express dystrophin (Fig. 2A and Supp. 8), transgenic myoblasts expressing WT-Dys exhibited lower VinTS BRET ratios (Fig. 2B) and therefore higher vinculin tension. As expected, VinTL BRET ratios were the same for NTg and WT-Dys myoblasts (Fig. 2C). As a control, we also characterized another transgenic myoblast line that expresses the non-pathogenic SNP in dystrophin, I232M. The tension measured at the I232M adhesion complexes using VinTS did not differ from WT-Dys myoblasts (Supp. 9A). These results show that stable dystrophin expression in myoblasts significantly impacts FA tension.

**Figure 2.**
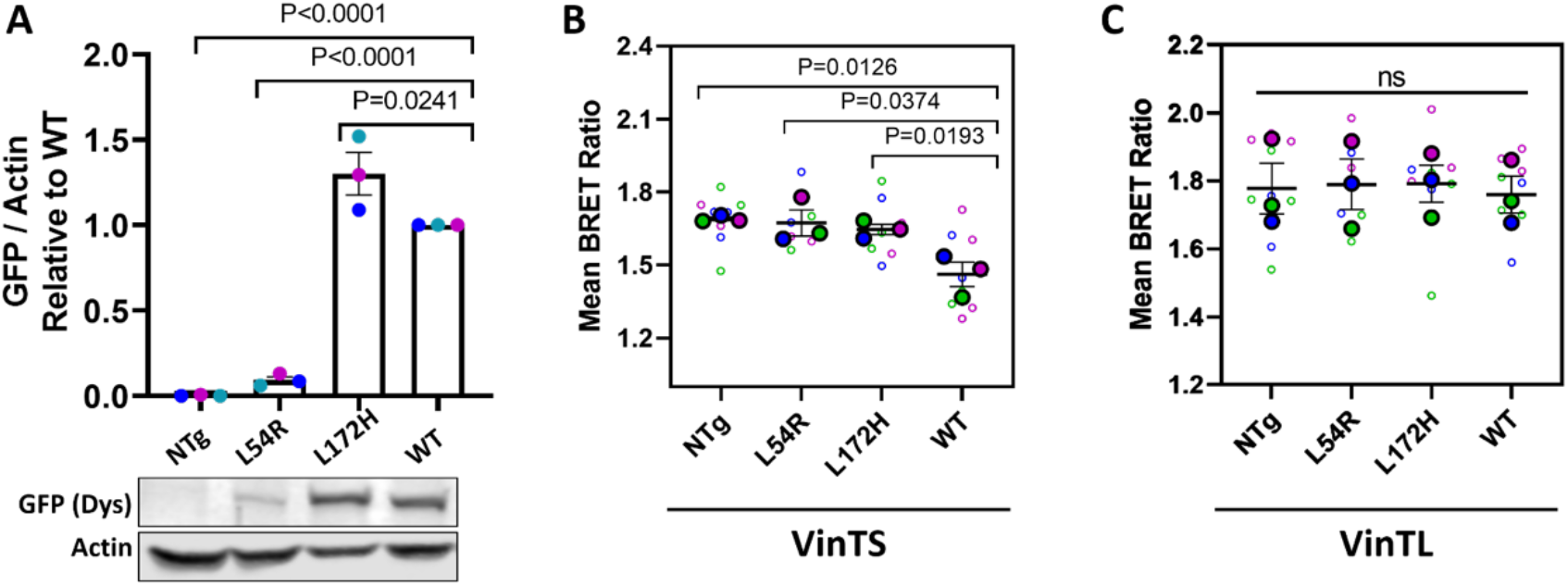
Dystrophin status influences focal adhesion tension. **A.** Quantification of N=3 separate cell lysates probed with GFP-dystrophin antibody (455 kDa) and normalized to Actin (42 kDa) (upper). Each color represents the same independent experiment set. Representative western blot of cell lysates probed for GFP-dystrophin and actin load control (lower). **B.** Apparent BRET efficiencies (N=3) for myoblasts transfected with VinTS and **C**. VinTL plated on fibronectin. Large dots denote the mean of independent experiments and smaller dots denote the individual mean BRET measurements per cell. Each experiment and corresponding individual measurements are colored the same. Western blot measurements are relative to its respective WT sample. Data analyzed via one-way ANOVA; ns, not significant. All error bars represent SEM.

### Dystrophy-associated mutations in dystrophin decrease focal adhesion tension

Missense mutations in dystrophin can lead to disease as severe as mutations causing complete loss of expression (Prior et al. 1993) through a pathomechanism that is not well understood. In order to test our hypothesis that forces between the extracellular matrix and cytoskeleton of muscle cells are disrupted by dystrophin mutations, we first characterized the transgenic cell lines expressing dystrophins encoding pathogenic L54R and L172H mutations, which map to actin binding domain 1. L54R is expressed at 8% of WT and L172H at 130%, (Fig. 2A) in myoblasts. As expected, non-load-bearing VinTL biosensors exhibit a relatively high BRET ratio, thus indicating low tension across all cell lines independent of dystrophin status (Fig. 2C).However, in contrast to the large decrease in BRET ratios observed in WT-Dys myoblasts, signifying higher tensions, the dystrophin missense mutant cells show relatively small decreases in BRET ratios more comparable to NTg (Fig. 2B and Supp. 10). These data suggest that the L54R and L172H mutations compromise the mechanical function of dystrophin.

Moreover, these data raise the question of whether the observed effects are due to functional disruptions in dystrophin that lead to pathogenicity versus altered dystrophin levels. As previously shown, I232M FA tension does not differ from WT-Dys expressing myoblasts even though it expresses 80% more dystrophin than WT-Dys myoblasts (Supp. 9B). This result shows that increased levels of dystrophin with a benign mutation in its actin binding domain do not affect FA tension. Thus, the decreased FA tension observed in myoblasts expressing L54R and L172H mutations suggests that the disease-causing missense mutations are modulating FA tension, although decreased levels of L54R dystrophin may also contribute to the reduction in measured tension.

We surveyed other select FA and DGC protein levels to rule out effects on vinculin tension. Paxillin, utrophin, α-dystroglycan and β-dystroglycan levels are the same for all dystrophin expressing cell lines (Supp. 11). Moreover, FA and cell morphology was the same for all studied cell lines (Supp. 12 and 13). These data indicate that dystrophin levels and/or missense mutations are the main cause for decreased tension transmitted along the FA load-bearing protein, vinculin.

### YAP mechanoactivation is altered by DMD- and BMD-associated mutations

Previous work in non-muscle cells has shown that altered FA tension correlates with disruption of mechanotransduction signaling pathways in the cell (Elosegui-Artola et al. 2017; Driscoll et al. 2015; Dasgupta and McCollum 2019). Thus, we investigated whether the function of a key mechanotransduction regulator YAP is altered in cell lines expressing mutant dystrophin. YAP is a transcriptional co-activator in many cell types, including myoblasts, and participates in maintaining homeostasis that is dysregulated in muscle disorders (Dupont et al. 2011; Fischer et al. 2016). FA tension generally promotes YAP activation (Dasgupta and McCollum 2019), which involves a change in localization from the cytoplasm to the nucleus. Thus, we first quantified nuclear/cytoplasmic ratios of YAP from fixed and immunostained myoblasts. Myoblasts expressing WT dystrophin showed an increased nuclear/cytoplasmic YAP ratio compared to NTg, and myoblasts expressing dystrophin mutations had comparable YAP ratios to NTg (Fig. 3A-B). Thus, WT-Dys myoblasts have increased YAP activation compared to myoblasts expressing DMD and BMD-associated mutations. In a separate set of experiments, we observed that I232M myoblasts YAP ratios showed no differences when compared to WT-Dys myoblasts (Supp. 9D-E). Our data suggests that in myoblasts, YAP is a mechanotransducer of dystrophin-mediated intracellular tension, which becomes dysregulated by expression of missense mutations in the actin binding domain 1 of dystrophin. The transcriptional co-activator with PDZ-binding motif (TAZ) is a paralog of YAP; changes in its activity were not a subject of this study.

**Figure 3.**
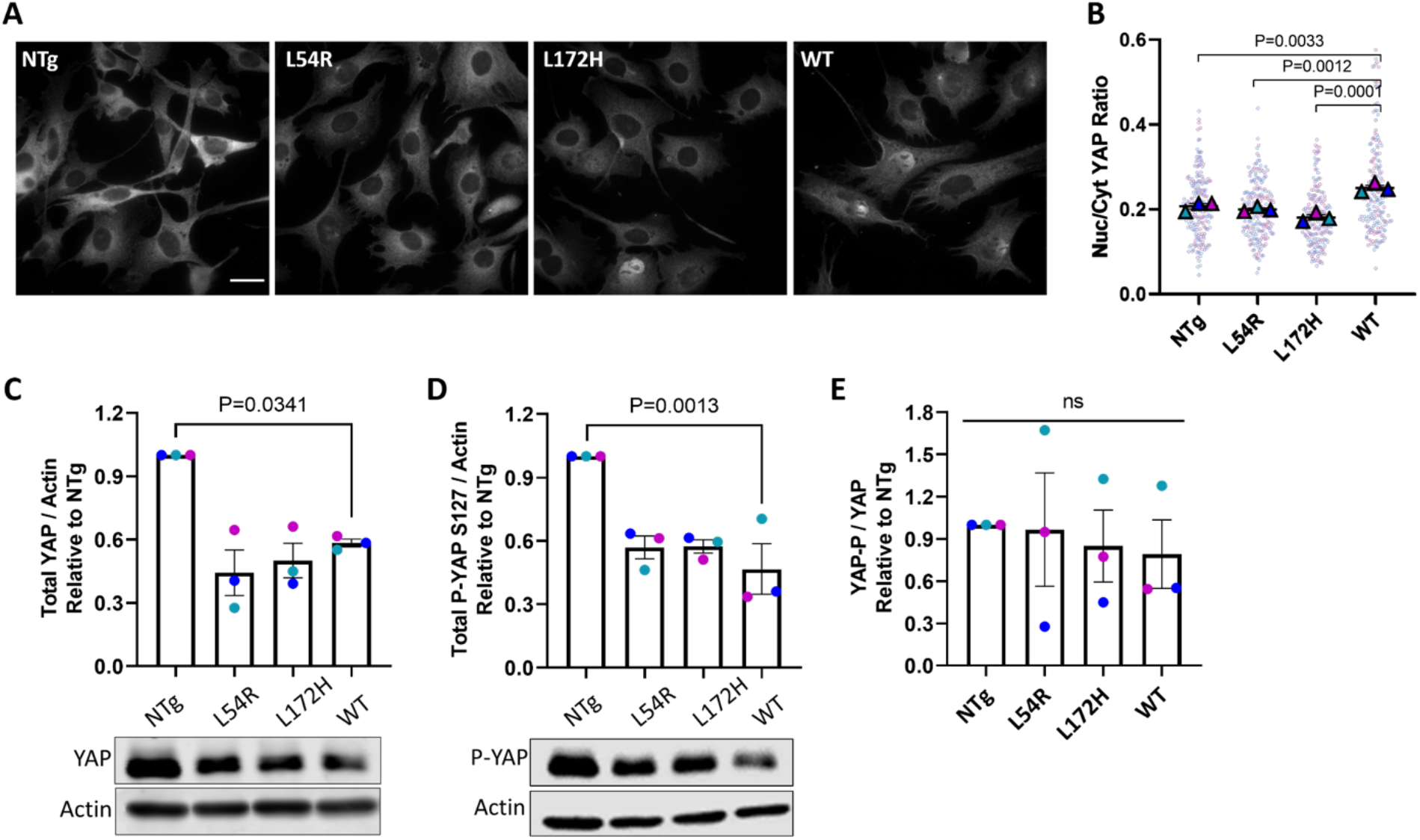
Dystrophin missense mutations alter YAP mechanoactivation. **A.** Representative images of fixed C2C12 myoblasts on fibronectin immunostained with YAP antibody. Scale bar 30 μm. **B.** Quantification of N=3 ratio of nuclear to cytoplasmic YAP. Large triangles denote the mean of independent experiments and smaller dots denote the mean of individual measurements. Each experiment and corresponding individual measurements are colored the same. **C.** Quantification of N=3 separate cell lysates probed with a YAP antibody (400 kDa) and a **D.** Ser 127 phospho-YAP antibody, normalized to Actin (42 kDa) (upper). Representative western blot of cell lysates probed for Utrophin and actin load control (lower). Western blot measurements are relative to its respective NTg sample. **E.** The ratio of the normalized P-YAP levels to the normalized YAP levels. Data analyzed via one-way ANOVA; ns, not significant. All error bars represent SEM.

Because YAP is a mechanosignaling node at the intersection of multiple signaling pathways (Ma et al. 2019), we interrogated how some of these pathways were affected by expression of WT and mutant dystrophins. The Hippo pathway has long been considered to be the canonical inhibitor of YAP activity through phosphorylation (Fischer et al. 2016). Phosphorylation of YAP on serine 127 (P-S127) via the large tumor suppressor kinase 1/2 (LATS1/2) downstream of the Hippo signaling cascade inhibits YAP activity through the binding of 14-3-3 proteins, leading to cytoplasmic sequestration and subsequent possible proteasomal degradation(Ma et al. 2019). Tension sensed and transmitted via FA is known to regulate YAP activity by various and complex pathways, dependent and independent of Hippo signaling, by activating the focal adhesion kinase (FAK). FAK can phosphorylate LATS 1/2 to inhibit its function or directly phosphorylate YAP on tyrosine residues to promote its nuclear translocation (Dasgupta and McCollum 2019). To determine if the Hippo pathway is involved in YAP cytoplasmic retention via P-S127, we analyzed lysates from all cell lines and western blotted for total YAP and YAP P-S127. We found that total YAP and P-S127 protein were significantly decreased in WT-Dys expressing cells when compared to NTg control, while L54R and L172H were expressed at the same levels of YAP as WT-Dys with only a modest, non-significant increase of P-S127 (Fig. 3C-D). The P-YAP/Total YAP ratio was the same for all lines (Fig. 3E). For I232M myoblasts, total YAP levels, P-S127 levels, and P-YAP/Total YAP ratios did not differ from WT-Dys (Supp.9F-H). These results demonstrate that the Hippo pathway does not play a major role in the regulation of YAP via phosphorylation of S127, indicating that it is the changes in vinculin tension at the FA level that likely modulate YAP inactivation for L54R and L172H expressing cells. Direct regulation of YAP via FA tension could be due to mechanoactivation of FAK that phosphorylates YAP on activating tyrosines.

### Defective migration in myoblasts expressing mutant dystrophins

YAP is known to promote migratory and proliferative behaviors (Fischer et al. 2016). Therefore, we measured cell migration and proliferation in myoblasts expressing WT and mutant dystrophins. We performed wound healing assays on confluent monolayers of cells plated on fibronectin and followed wound closure (area free of cells) over time (Fig. 4A). The NTg confluent cell sheet migrated the fastest compared to all transgenic cell lines (Fig. 4B) despite its similar nuclear to cytoplasmic YAP ratios to L54R and L172H myoblasts. This may be explained by its increased levels of total YAP (Fig. 3C). The wound closure rates of the other cell lines better correlate with the nuclear to cytoplasmic YAP ratios. The collective cell sheet formed by WT-Dys and I232M cells filled the wound area faster than L172H and L54R cell sheets (Fig. 4B). These results show that the expression of dystrophin in myoblasts slows down their collective migration, regardless of the levels and/or mutations of dystrophin. Additionally, dystrophin missense mutations lead to further decrease of collective cell migration. Using the calculated wound closure rates (Fig. 4B), we inferred the average edge “speed” of the cell sheet front, to take into account the differences in initial wound size. The apparent edge “speeds’’ followed the same trend as the wound closure rates (Fig. 4C). Compared to NTg myoblasts, WT-Dys and I232M wound edge was slower by 50% and 69%, while compared to WT-Dys L172H and L54R were slower by 4% and 34%, respectively.

**Figure 4.**
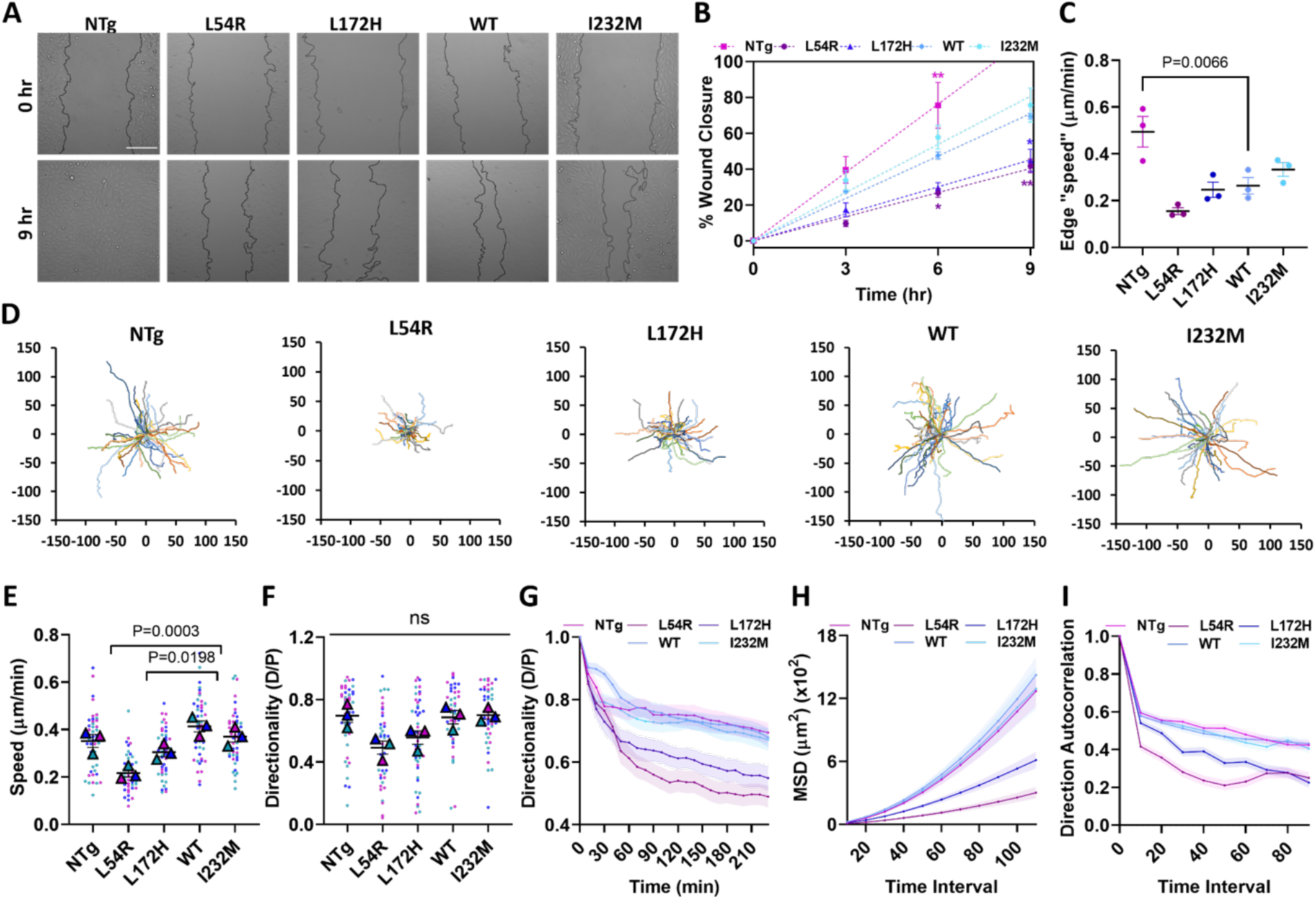
Missense mutations cause myoblasts migration defects. **A.** Representative images of wound-healing assays at 0 and 9hrs after scratching a confluent monolayer of cells cultured on fibronectin. Scale bar 200 μm. **B.** Comparison of percentage of wound closure over time for N=3 for each cell line. Data points fitted with a linear regression. **C.** Inferred edge speed of cell sheet calculated from the closure rates obtained from each linear regression fit (N=3). **D.** Origin plots for all cell trajectories (N=3, 15 trajectories each) of single migrating cells, from which **E.** cell speed, **F.** directionality ratio measured at the last point of trajectory, **G.** directionality ratio over time, **H.** mean square displacement and **I.** direction autocorrelation, were calculated to characterize cell migration efficiency. Large triangles denote the mean of independent experiments and smaller dots denote the mean of individual measurements per cell. Each experiment and corresponding individual measurements are colored the same. Data analyzed via one-way ANOVA; ns, not significant. All error bars represent SEM.

To gain more insight into the 2D migratory behavior of our myoblast lines, we cultured cells at low density on fibronectin to allow for the tracking of single cells. We imaged them over a period of 4 hrs. Qualitatively, we observed that WT-Dys, and I232M myoblast migration was comparable to NTg, and migrated more quickly and more directionally than L54R and L172H myoblasts, as shown by the migration tracks shown in figure 4D. Quantitatively, WT-Dys and I232M myoblasts all had comparable single cell speeds to NTg, while L172H and L54R myoblasts were 27% and 46% slower, respectively, when compared to WT (Fig. 4E and Supp. 14-15). However, directional persistence was not significantly different for any of the cell lines, even when there was correlation with cell speeds (Fig. 4F). Directionality was calculated by cell displacement over their path between the first and last point of trajectory. To highlight more nuanced differences in directional persistence between cell lines, we calculated directionality discretely over time rather than only at the last point. Doing so revealed greater differences between the directionality of cells expressing dystrophin missense mutants and the other cell lines. NTg, WT-Dys and I232M myoblasts directionality over time was the same, while L54R and L172H were significantly lower, each becoming progressively less directional over time (Fig. 4G). While progressive loss of directionality also occurs for NTg, WT, and I232M, it is less pronounced.

We also calculated the mean square displacements (MSD) as another approach to estimate persistence in order to gauge the area explored by the cell while migrating over increasing time intervals. As per our directionality ratio results, L54R and L172H myoblasts explore less area over time than NTg, WT, and I232M all of which had comparable values amongst each other (Fig 4H). Recent work by (Gorelik and Gautreau 2014) proposes to use direction autocorrelation to measure directional persistence, because directionality is biased by cell speed(Gorelik and Gautreau 2014). Using their open source program DiPer, we calculated direction autocorrelation as a function of time and observed the same trend for directionality but with more pronounced differences; the angles describing the trajectory of L54R and L172H cells are less aligned with each other over different time scales in comparison to NTg, WT, and I232M (Fig. 4I). MSD and direction autocorrelation were equivalent between L54R and L172H for almost all time intervals. These results demonstrate that missense dystrophin mutations associated with DMD and BMD cause myoblasts to migrate less efficiently, whereas not expressing dystrophin or expressing a benign SNP does not influence migration speed or directionality. Furthermore, these results highlight the pathogenicity of DMD-associated mutations. Although L54R myoblasts express 92% less dystrophin than WT expressing myoblasts, the low levels of mutant dystrophin still severely impairs migration.

We next measured cell proliferation in the cell lines to rule out potential confounding effects (Morikawa et al. 2017) on migration caused by increased nuclear YAP. However, we measured no differences between our cell lines in proliferation over four days (Supp. 16).

Altogether, decreased FA tension, YAP inactivation, and slower migration suggest that disease-associated missense mutations in dystrophin disrupt a link between tension transmission and YAP mechanoactivation in myoblasts.

## Discussion

Force-transmitting complexes from the membrane to the nucleus allow for the transduction of physical cues into biochemical signals and the regulation of gene expression. In disease, the mechano-responsiveness of these complexes and their connection to different elements of the mechanotransduction axis can be altered. A phenotype of muscle cells harboring abnormal dystrophin is a destabilized sarcolemma due to weaker cytoskeletal-ECM coupling (Lapidos, Kakkar, and McNally 2004), that leads to membrane breakage and increased compliance (Menke and Jockusch 1991; Mokri and Engel 1998; García-Pelagio et al. 2011; Lopez et al. 2021). Remarkably, we show here that this phenotype manifests even at a molecular level as decreased focal adhesion tension, which disrupts mechanotransduction pathways in the cell. Thus, we propose that disease-associated missense mutations in actin binding domain 1 compromise dystrophin’s role as a critical mechanical link between the ECM and cytoskeleton.

An intriguing observation was the absence of tension gradients along the length of single FA for VinTS (Supp. 7-8), which has been previously observed by us and others (Sarangi et al. 2017; Rothenberg et al. 2018; Aird et al. 2020). The lack of tension gradients could be due to the cell type and differentiation stage. In the context of mature skeletal muscle, myoblasts are activated satellite cells. Upon muscle injury, satellite cells activate into myoblasts and crawl over the basal lamina of the fiber, which is composed primarily of collagen IV and laminin, until they reach the injury site. Interstitial fibroblasts initially coat the site with a fibrin, tenascin, and a fibronectin scaffold, where myoblasts align and differentiate into myotubes to repair the wound (Goetsch, Myburgh, and Niesler 2013). Our experiments were performed on myoblasts cultured on fibronectin, an environment that after muscle injury signals the cell to remain and fuse to the surrounding fibers. Therefore, it is possible that due to the nature of the fibronectin-myoblast interaction, the cells homogeneously distribute their tension at the FA to prepare for subsequent fusion, similar to how they would fuse in mature injured skeletal muscle.

There are conflicting reports on the role and regulation of YAP in skeletal muscle homeostasis and disease (Fischer et al. 2016). YAP protein levels and activation reported in muscular dystrophy are varied, showing increased, decreased, or equal levels when compared to healthy controls (Hulmi et al. 2013; Vita et al. 2018; Iyer et al. 2019). In cardiomyocytes, the Hippo pathway appears to play a significant role in the regulation of YAP with direct involvement of the DGC. Lack of dystrophin in *mdx* mice caused dysregulation of Hippo signaling with direct involvement of the DGC, leading to cardiomyocyte overproliferation after injury (Morikawa et al. 2017). However, Morikawa et al., also showed using myotubes differentiated from C2C12 myoblasts that dystroglycan binds to and directly sequesters YAP near the sarcolemma, while knocking down the *Dmd* gene led to decreased YAP localization at the sarcolemma and cytoplasm, and increased YAP nuclear localization. Proliferation dysregulation was not explored for C2C12 myotubes, but their *mdx* and myotube data was used to reach their final conclusion. In contrast, we showed that myoblasts expressing DMD- and BMD-associated missense mutations exhibited decreased YAP activation (Fig. 3A-B), no involvement of the Hippo pathway (Fig. 3C-E), and equal proliferation to NTg, WT and I232M lines (Supp. 16). A major factor that could account for these differences is that their *Dmd* knockdown cells do not necessarily reflect a pathogenic state. Cultured myoblasts or fibers, as used in their study, do not express dystrophin to begin with, while our transgenic lines do express dystrophin, both healthy and associated with muscular dystrophies. Nonetheless, as highlighted these conflicting reports on phosphorylation and activation may be due to several factors such as the muscle sample source, the type of muscle, harvest time, and the cellular differentiation stage.

Additionally, YAP phosphorylation increases during muscle differentiation while YAP expression declines (Fischer et al. 2016). In skeletal muscle, dystrophin expresses and completely localizes to the sarcolemma in the later stages of cell differentiation and embryonic development (Chevron et al. 1994). NTg myoblasts express higher levels of YAP compared to all transgenic lines (Fig. 3C), suggesting that transgenic expression of dystrophin, healthy or diseased, could mimic a more advanced stage of cell differentiation in regards to dystrophin and possible maturation of the DGC. Our data shows that NTg cells have decreased FA tension (Fig. 2A), a decreased nuclear/cytoplasmic YAP ratios (Fig. 3A-B), and comparable single cell migration rates and directionality (Fig. 4D-I) when compared to WT-Dys myoblasts. A possible explanation for this is that healthy – albeit dystrophin-less – NTg cultured myoblasts compensate for this lessened tension during their early development by overexpressing YAP until they begin to express dystrophin in the later stages of differentiation. Whether this is the case in developing muscle remains to be determined.

A surprising result was that L54R and L172H myoblasts migrate more slowly than WT-Dys and I232M, even in light of our YAP results. As previously explained, dystrophin is expressed and localized to the sarcolemma later during the development to reinforce the sarcolemma and to protect it against mechanical stress, when myoblasts have already become fused fibers. Moreover, once it localizes to the subsarcolemma, it becomes part of the DGC, essentially becoming a specialized FA in skeletal muscle, which gives it increased adhesion contacts to the ECM. Therefore, we expected cells expressing WT and I232M dystrophin to migrate more slowly than cells expressing mutant defective dystrophins, but our results reveal the opposite. Analysis from patient muscles show decreased levels of L54R and L172H mutant dystrophin, although they correctly localize to the sarcolemma (Prior et al. 1993; Hamed et al. 2005). These findings suggest that the remaining dystrophin is less functional and perhaps “poisons’’ the cell. Both missense mutations are in the actin binding domain of dystrophin. We have previously shown *in vitro* that L54R mutation causes only a 4-fold decrease in actin binding affinity while L172H had no effect, yet both dystrophin mutants were less stable compared to WT *in vitro* (Henderson, Lee, and Ervasti 2010) and *in vivo* (Talsness, Belanto, and Ervasti 2015). The unifying factor between our molecular tension, YAP activation and cell migration data is the actin cytoskeleton. Although our previous actin cosedimentation data did not show any major differences regarding actin affinity of dystrophin, our migration data suggests that actin dynamics may be impaired in myoblasts expressing L54R and L172H dystrophin mutants. Migration is driven by the continuous reorganization of the actin cytoskeleton, and it depends on a complex interplay between the cytoskeleton and several signaling pathways (Choi, Ferrari, and Tedesco 2020). Future studies will determine how the dynamic interactions between actin and dystrophin affect migration in muscle regeneration and disease.

YAP activity promotes tissue regeneration and repair, and muscle regeneration is impaired in muscular dystrophy (Cai et al. 2010; Hulmi et al. 2013; Xin et al. 2013; Wallace and McNally 2009). Here we demonstrate that DMD- and BMD-associated dystrophin mutations lead to migration defects in myoblasts, and migration is critical for regeneration to occur. Over time, dystrophic muscle loses its regenerative potential, but there is no consensus on how this regenerative exhaustion occurs. Our results suggest that impaired cell migration may contribute to the decreased regenerative capacity of dystrophic muscle caused by missense mutations in the actin binding domain.

## ACKNOWLEDGEMENTS

Authors thank Karen Evans for manuscript revision and Polina Alexandrovich for assistance with data compilation and Yahor Savich for helpful discussion. Authors would like to acknowledge funding from NIH (R01AR042423, R01AR049899, and R35GM119483) and the Muscular Dystrophy Association. WRG is a Pew Biomedical Scholar. MR would like to acknowledge funding from the University of Minnesota Frieda M. Kunze and Doctoral Dissertation Fellowships.

