## Supplementary info for "Dystrophin modulates focal adhesion tension and YAP-mediated mechanotransduction"

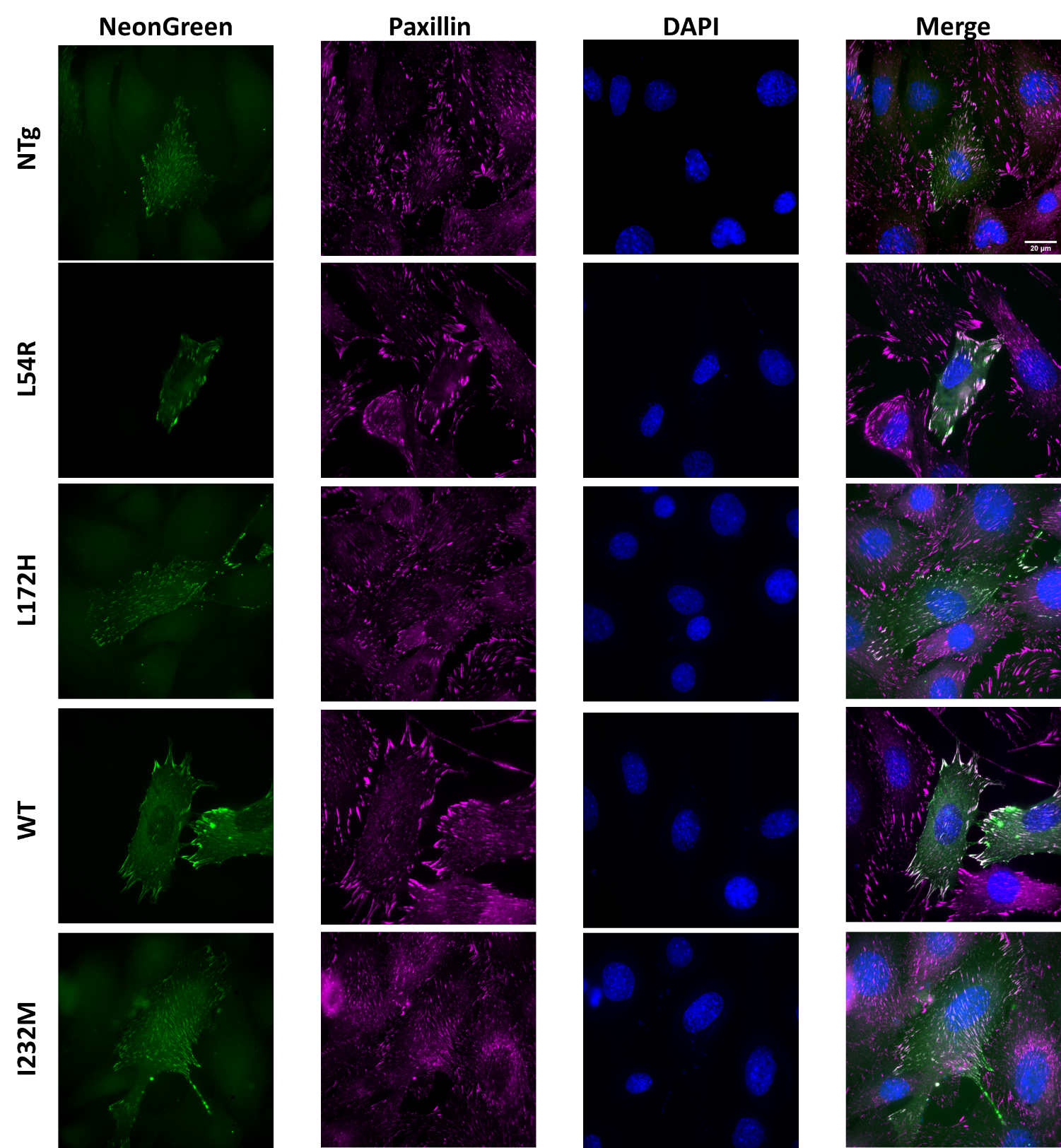

**Supplementary Figure 1. VinTS localizes to focal adhesions.** Representative image of fixed C2C12 myoblasts transfected with VinTS (NeonGreen; NG), immunostained with paxillin (magenta) and DAPI (blue) to indicate focal adhesions and nucleus, respectively. Merged images show co-localization of the tension sensor with paxillin.

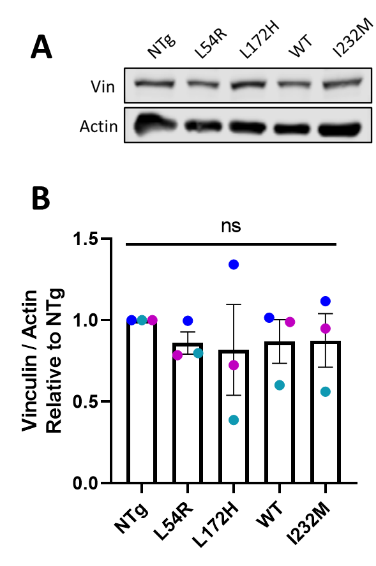

**Supplementary Figure 2. Vinculin expression levels. A.** Representative western blot of cell lysates probed for vinculin (116 kDa) and actin (42 kDa) as a load control. **B.** Quantification of vinculin levels of N=3 separate transfected myoblasts cell lysates. Each circle color represents the same independent experiment set. Western blot measurements are relative to its respective NTg control sample. Data analyzed via one-way ANOVA; ns, not significant. All error bars represent SEM.

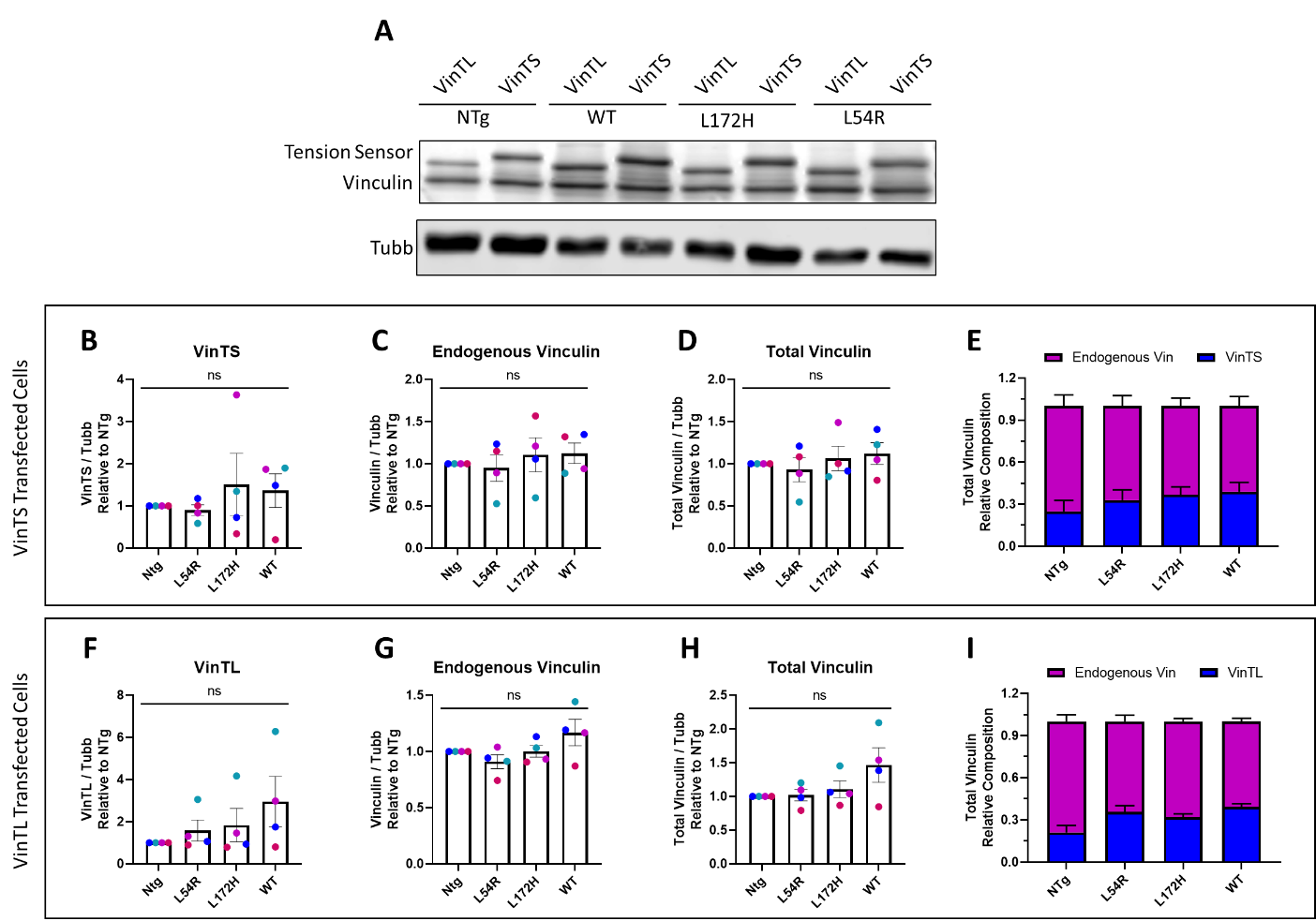

**Supplementary Figure 3. Vinculin and tension sensors expression levels across cell lines. A.** Representative western blot of cell lysates probed with a vinculin antibody and alpha tubulin as a load control. VinTS is 143 kDa, VinTL 164 kDa, vinculin is 116 kDa and alpha tubulin is 55 kDa. **B.** Quantification of VinTS and **C.** Vinculin expression levels of N=4 separate transfected myoblasts cell lysates. **D.** Sum of VinTS and endogenous vinculin to compare total vinculin cell levels across all myoblast lines. **E.** Proportion of VinTS and endogenous vinculin from total vinculin content. **F.** Quantification of VinTL and **G.** Vinculin expression levels of N=4 separate transfected myoblasts cell lysates. **H.** Sum of VinTL and endogenous vinculin to compare total vinculin cell levels across all myoblast lines. **I.** Proportion of VinTL and endogenous vinculin from total vinculin content. Each circle color represents the same independent experiment set. Western blot measurements are relative to its respective NTg control sample. Data analyzed via one-way ANOVA; ns, not significant. All error bars represent SEM.

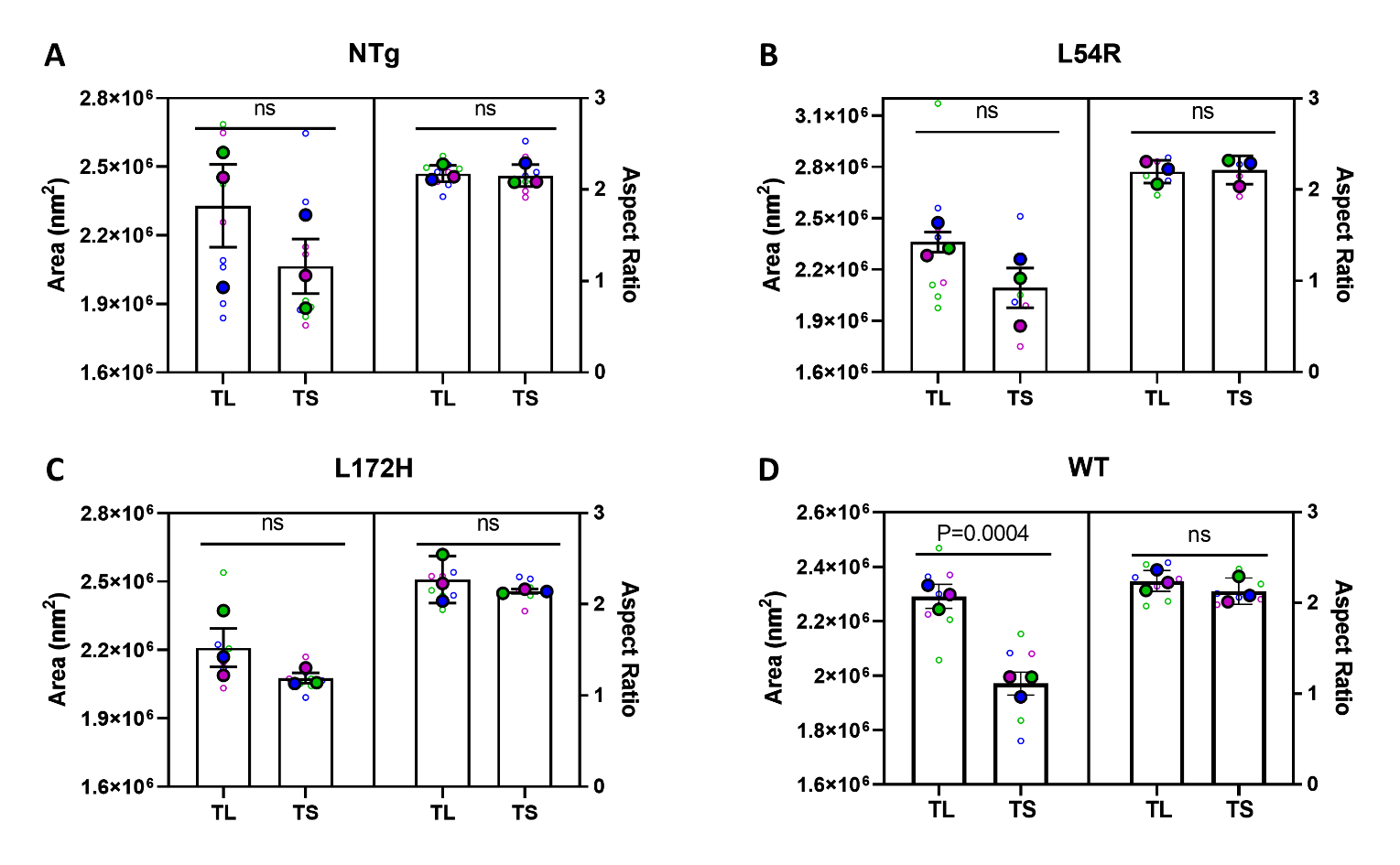

**Supplementary Figure 5. VinTS vs VinTL focal adhesion morphology**. Focal adhesion morphology comparison (N=3) of **A-D.** NTg, L54R, L172H and WT myoblasts transfected with VinTL and VinTS, respectively. Morphology parameters compared are area and aspect ratio. Large dots denote the mean of independent experiments and smaller dots denote the mean of individual focal adhesion measurements per cell. Each experiment and corresponding individual measurements are colored the same. Data analyzed via an unpaired 2-tailed t-test; ns, not significant. All error bars represent SEM.

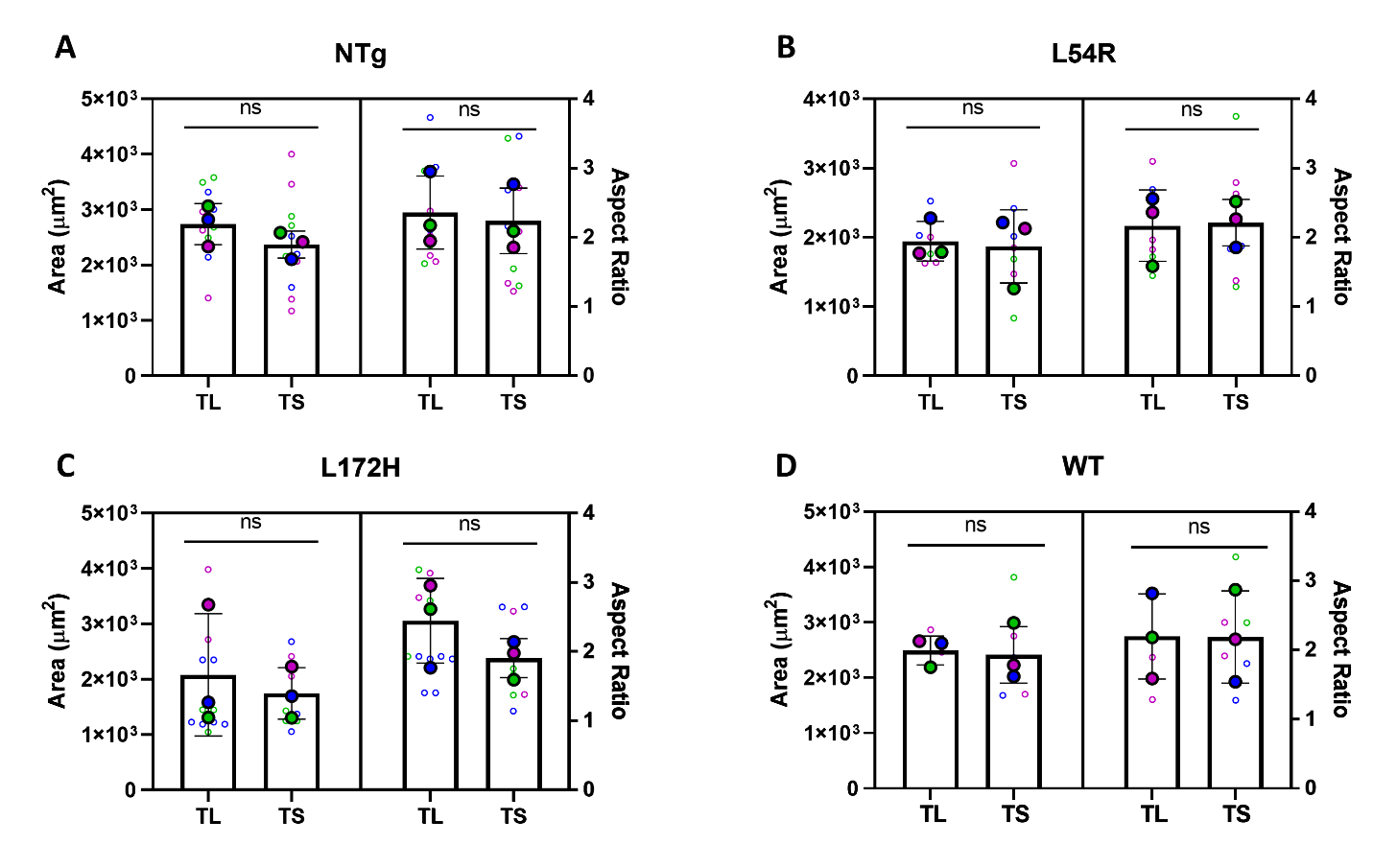

**Supplementary Figure 5. VinTS vs VinTL cell morphology.** Cell morphology comparison (N=3) of **A-D.** NTg, L54R, L172H and WT myoblasts transfected with VinTL an VinTS, respectively. Morphology parameters compared are area and aspect ratio. Large dots denote the mean of independent experiments and smaller dots denote the mean of individual measurements. Each experiment and corresponding individual measurements are colored the same. Data analyzed via an unpaired 2-tailed t-test; ns, not significant. All error bars represent SEM.

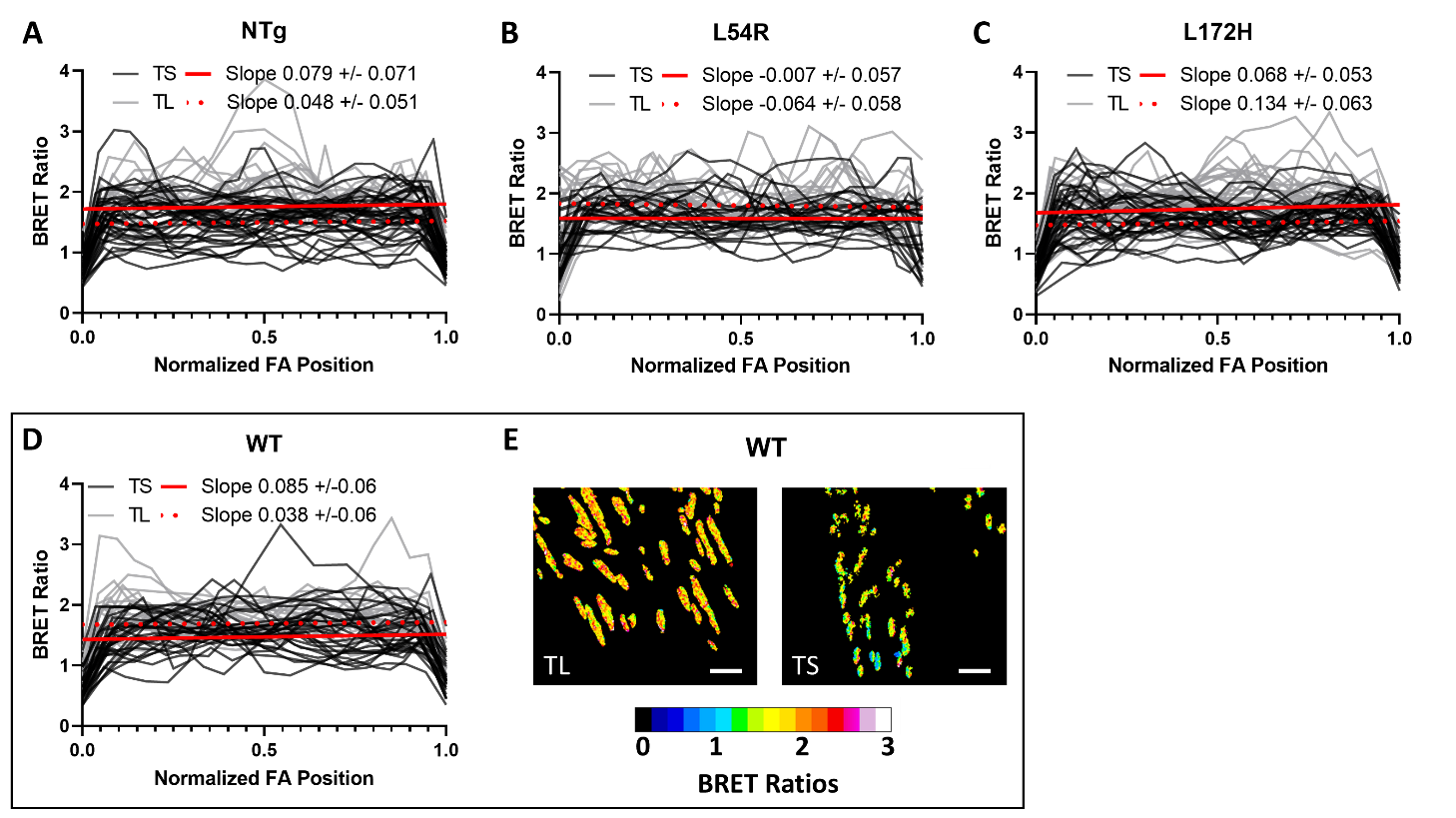

**Supplementary Figure 6. VinTS vs VinTL tension gradients**. **A-D.** Manual line scans along VinTS (black) and VinTL (grey) focal adhesions, normalized to the focal adhesion (FA) length (24-36 total focal adhesions from N=3 independent experiments). Shown in red, the linear regression of all line scans combined for VinTS (solid line) and VinTL (dotted line). The linear regression slope is given ± the standard deviation of the linear regression calculation. **E.**  Processed ratiometric images of focal adhesions of Dys-WT myoblasts. Image scale bar is 3 μm, color scale bar displayed as BRET ratio.

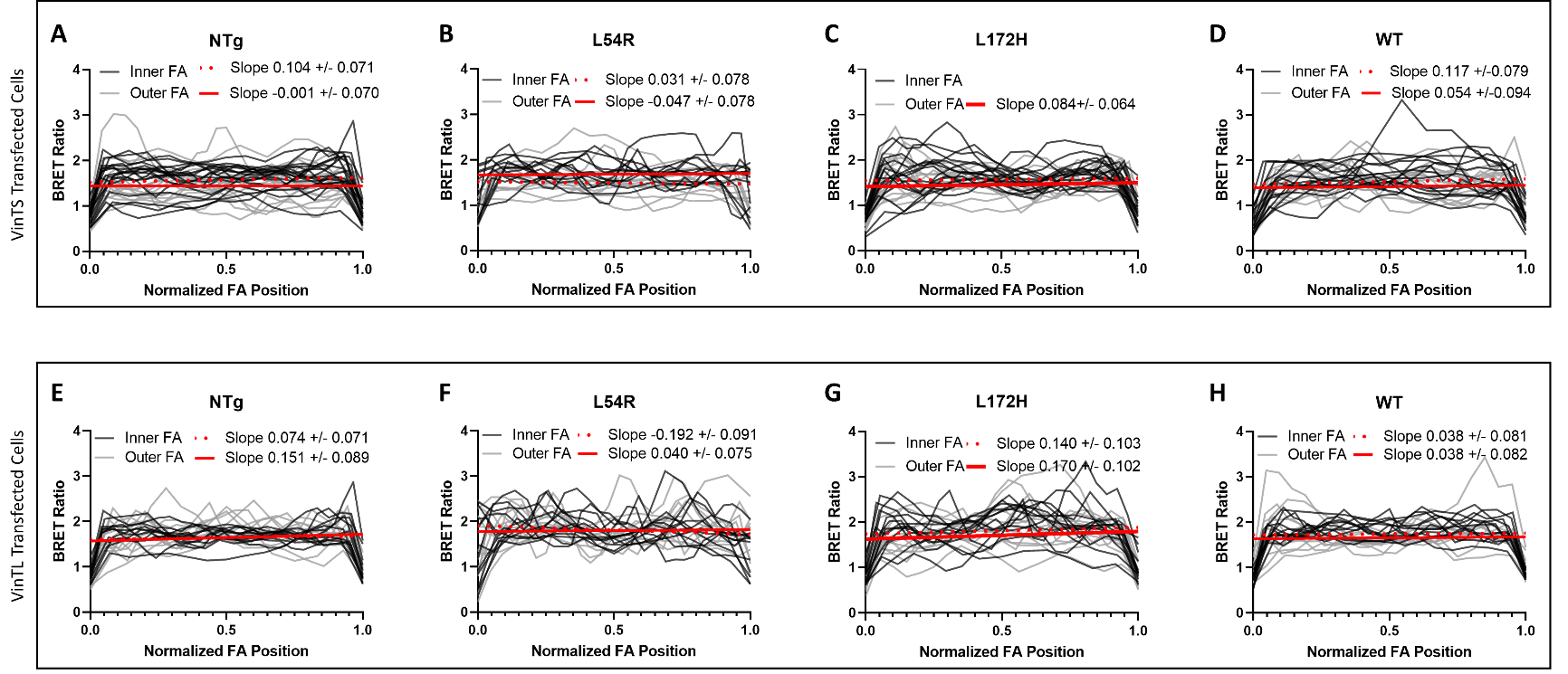

**Supplementary Figure 7. Inner vs outer focal adhesion tension gradients**. **A-D.** Manual line scans along VinTS and **E-H.** VinTL from inner (black) and outer (grey) focal adhesions, normalized to the focal adhesion (FA) length (12-18 total focal adhesions from N=3 independent experiments). Shown in red, the linear regression of all line scans combined for inner (dotted line) and outer (solid line). The linear regression slope is given ± the standard deviation of the linear regression calculation.

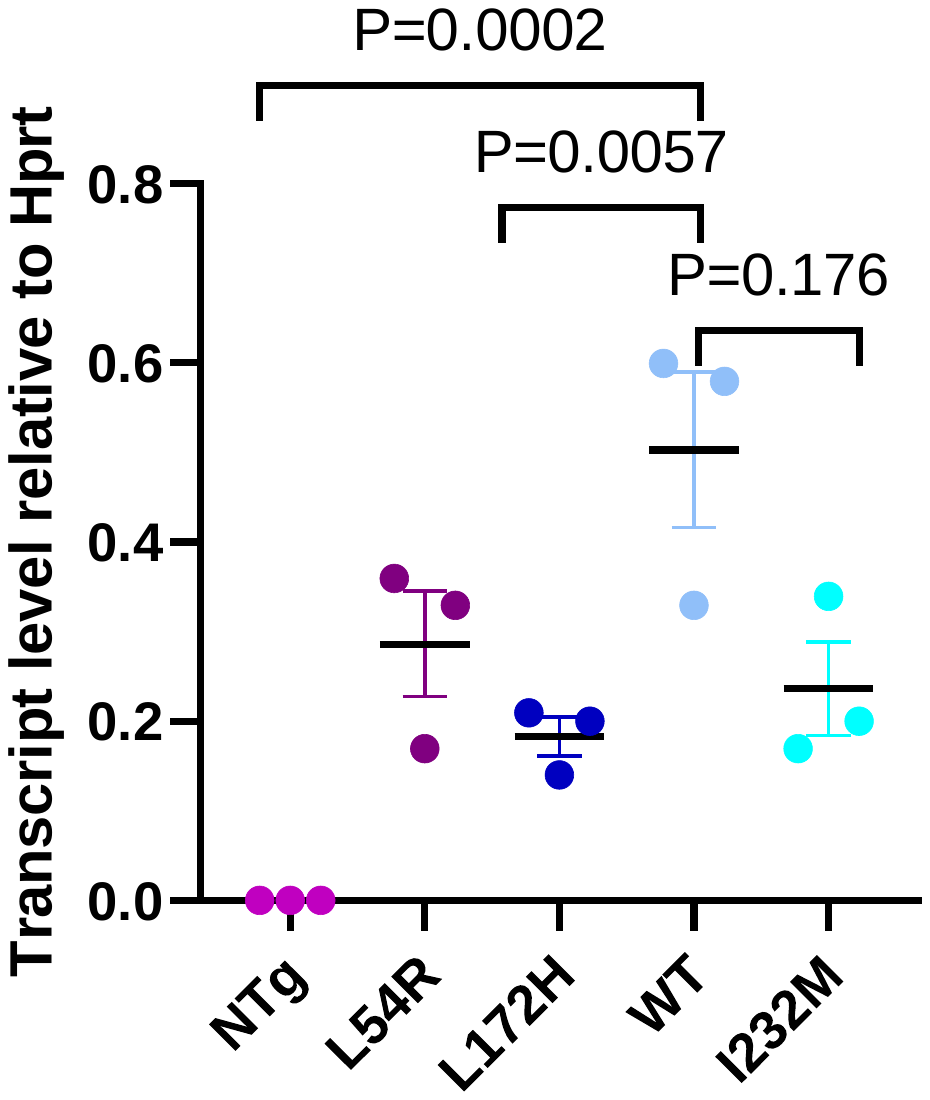

**Supplementary Figure 8. RT-qPCR of GFP-Dystrophin RNA.** Quantification of N=3 independent experiments measuring GFP-Dystrophin transgenic transcript levels relative to reference transcript Hprt. Data analyzed via one-way ANOVA; ns, not significant. All error bars represent SEM.

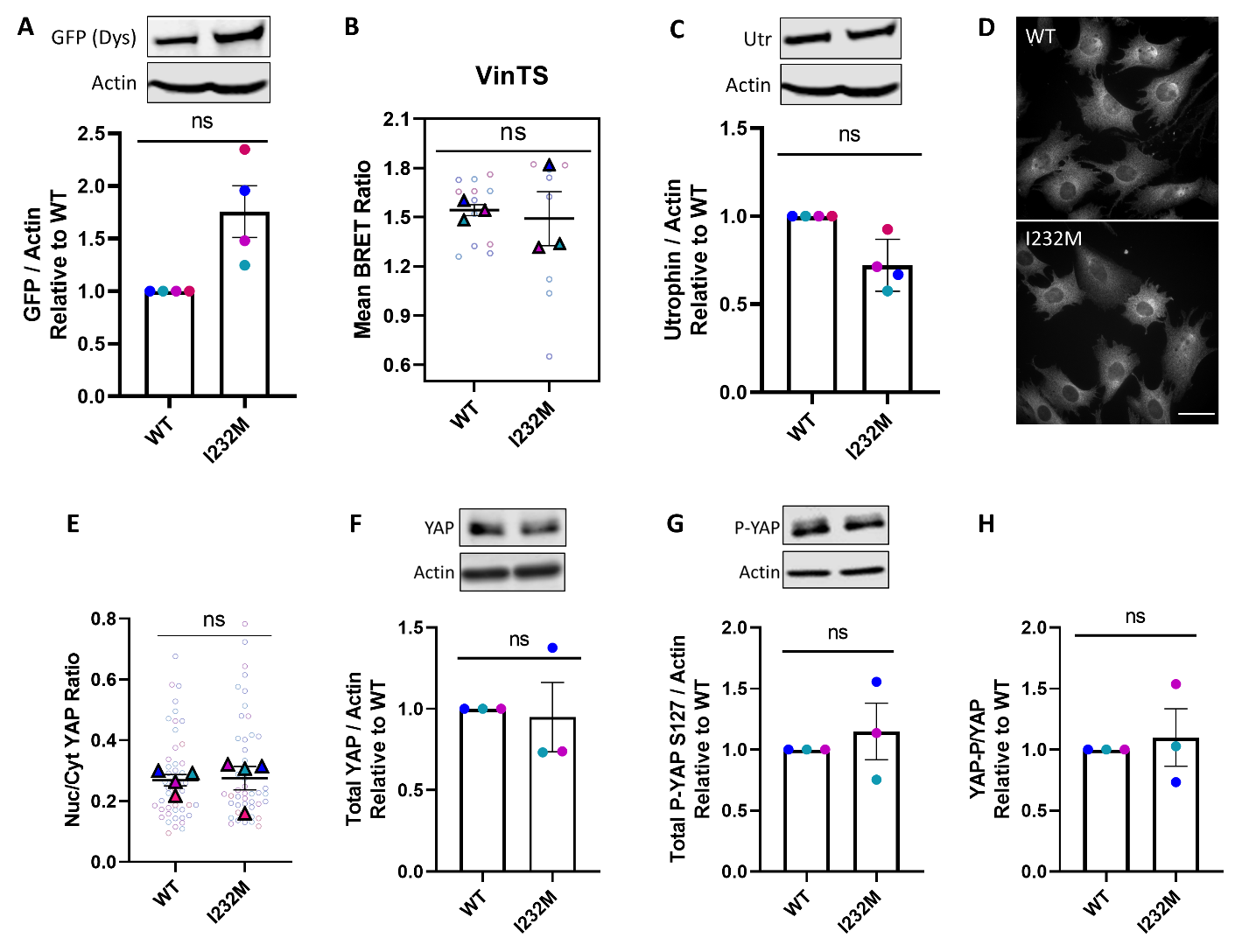

**Supplementary Figure 9. I232M has the same focal adhesion tension and YAP activation as WT myoblasts. A.** Representative western blot of cell lysates probed for GFP-dystrophin and actin load control (upper). Quantification of N=4 separate cell lysates probed with GFP-dystrophin antibody (455 kDa) and normalized to Actin (42 kDa) (lower). Each color represents the same independent experiment set. **B.** Apparent BRET efficiencies (N=3) for myoblasts transfected with VinTS. Large triangles denote the mean of independent experiments and smaller dots denote the individual mean BRET measurements per cell. Each experiment and corresponding individual measurements are colored the same. **C.** Representative western blot of cell lysates probed for Utrophin and actin load control (upper). Quantification of N=4 separate cell lysates probed with Utrophin antibody (400 kDa) normalized to Actin (42 kDa) (lower). **D.** Representative images of fixed C2C12 myoblasts on fibronectin immunostained with YAP antibody. Scale bar 40 μm. **E.** Quantification of N=4 ratio of cytoplasmic over nuclear YAP. Large triangles denote the mean of independent experiments and smaller dots denote the mean of individual measurements. Each experiment and corresponding individual measurements are colored the same. **F.** Quantification of N=3 separate cell lysates probed with a YAP antibody (400 kDa) and a **G.** Ser 127 phospho-YAP antibody, normalized to Actin (42 kDa) (upper). **H.** The ratio of the normalized P-YAP levels to the normalized YAP levels. Western blot measurements are relative to its respective WT sample. Data analyzed via an unpaired 2-tailed t-test; ns, not significant. All error bars represent SEM.

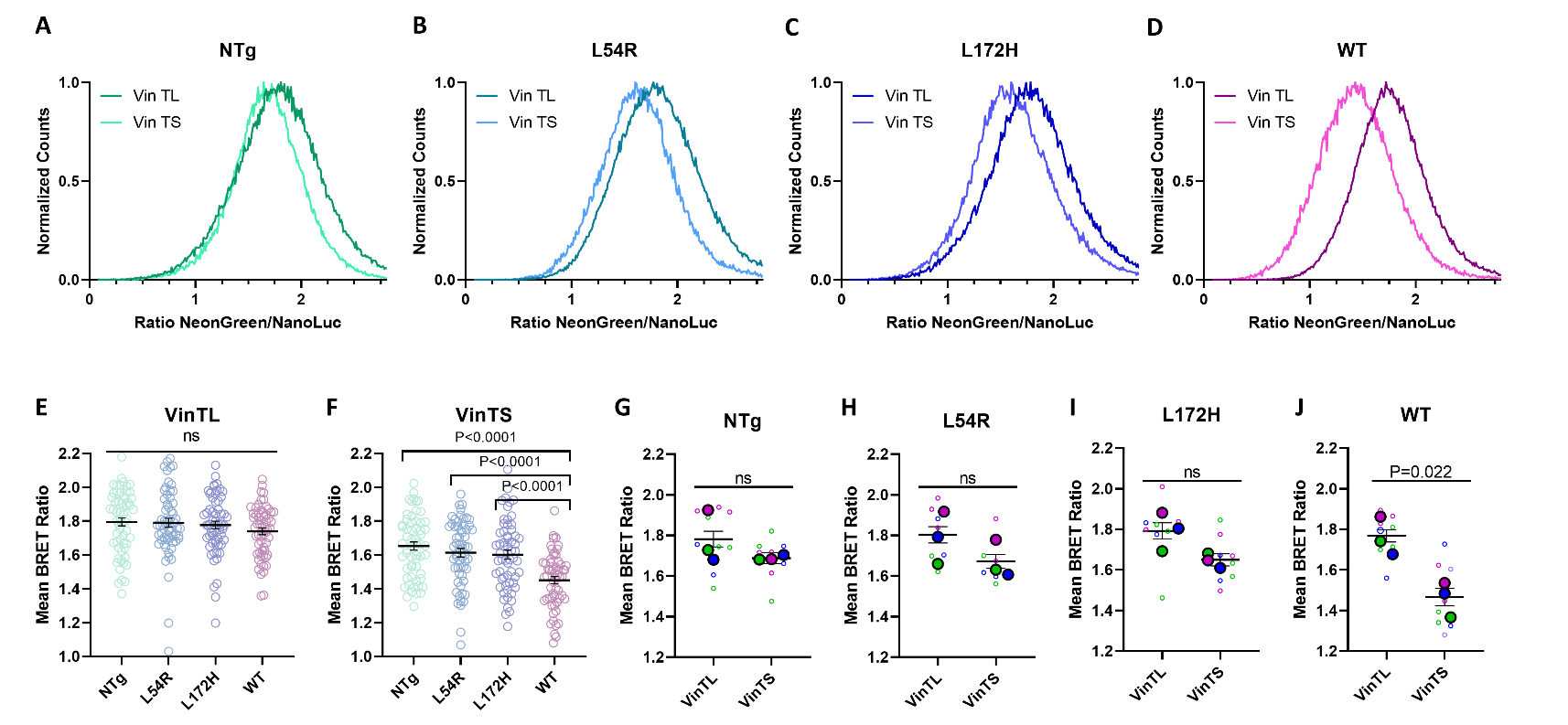
**Supplementary Figure 10.**  **BRET ratio comparison of VinTS vs VinTL across cell lines. A-D.** Normalized distribution of BRET ratios, **E-F.** Mean BRET ratios of single focal adhesions (60 total focal adhesions from N=3 independent experiments) and **G-J.** Calculated distribution mean for each individual distribution for VinTS and VinTL transfected myoblasts NTg, L54R, L172H and WT, respectively (N=3). Large dots denote the mean of independent experiments and smaller dots denote the individual mean BRET measurements per cell. Each experiment and corresponding individual measurements are colored the same. E-F data analyzed via one-way ANOVA and G-J data analyzed via an unpaired 2-tailed t-test; ns, not significant. All error bars represent SEM.

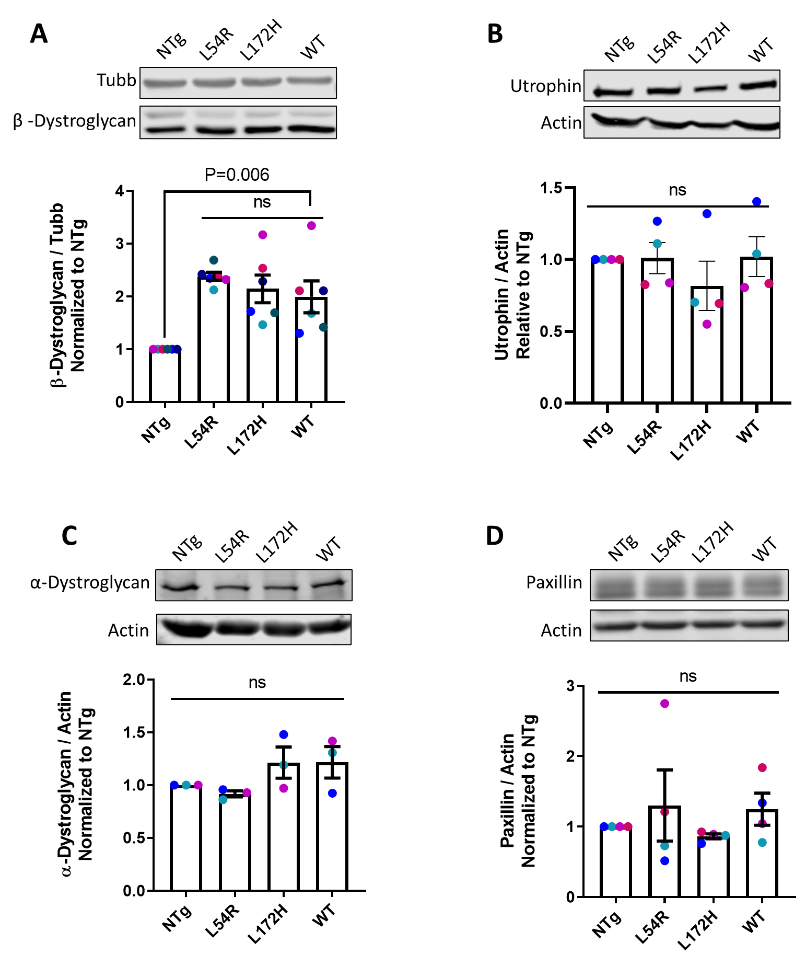

**Supplementary Figure 11. Dystrophin-glycoprotein complex and focal adhesion protein levels.** Representative western blot of cell lysates (upper) and quantification of separate transfected myoblasts lysates (lower), probed with **A.** Beta-dystroglycan, 43 kDa (N=6) **B.** Utrophin, 400 kDa (N=4) **C.** Alpha-dystroglycan, 156 kDa (N=3) and **D.** Paxillin, 65 kDa (N=4) antibodies, with their respective load controls. Each circle color represents the same independent experiment set. Western blot measurements are relative to its respective NTg control sample. Data analyzed via one-way ANOVA; ns, not significant. All error bars represent SEM.

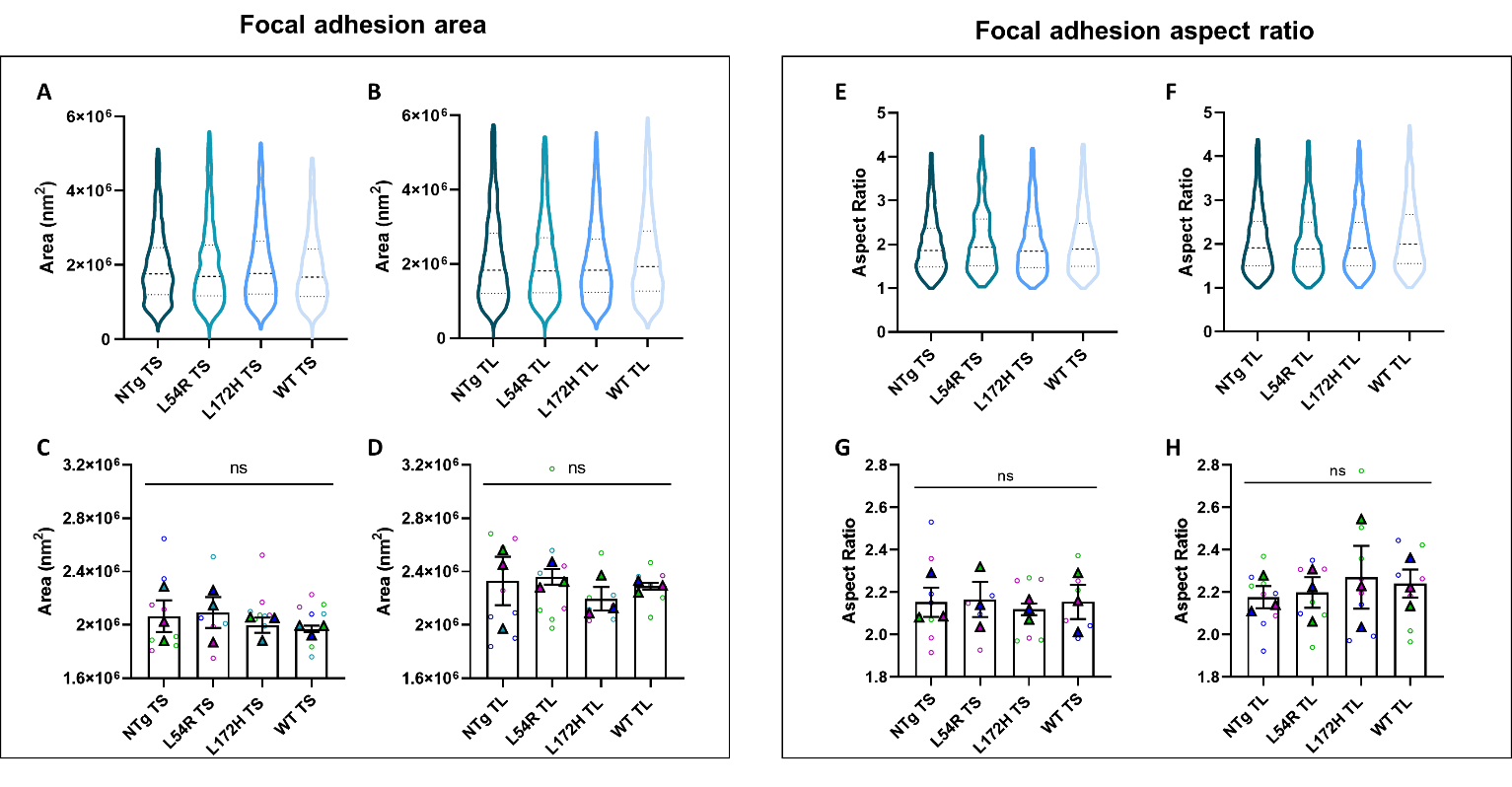

**Supplementary Figure 12. Focal adhesion morphology across cell lines.** Violin plots of area comparison for **A.** VinTS and **B.** VinTL transfected myoblast lines, for all individual focal adhesions data (>1000 data points). Mean focal adhesion area for N=3 independent experiments (large triangles) and mean of individual focal adhesion area per cell (small dots) for **C.** VinTS and **D.** VinTL. Violin plots of aspect ratio comparison for **E.** VinTS and **F.** VinTL cells, for all individual focal adhesions data. Mean aspect ratio data for N=3 independent experiments (large dots) and mean of individual focal adhesion aspect ratio per cell (small dots) for **G.** VinTS and **H.** VinTL. Each experiment and corresponding individual measurements are colored the same. Data analyzed via one-way ANOVA; ns, not significant. All error bars represent SEM.

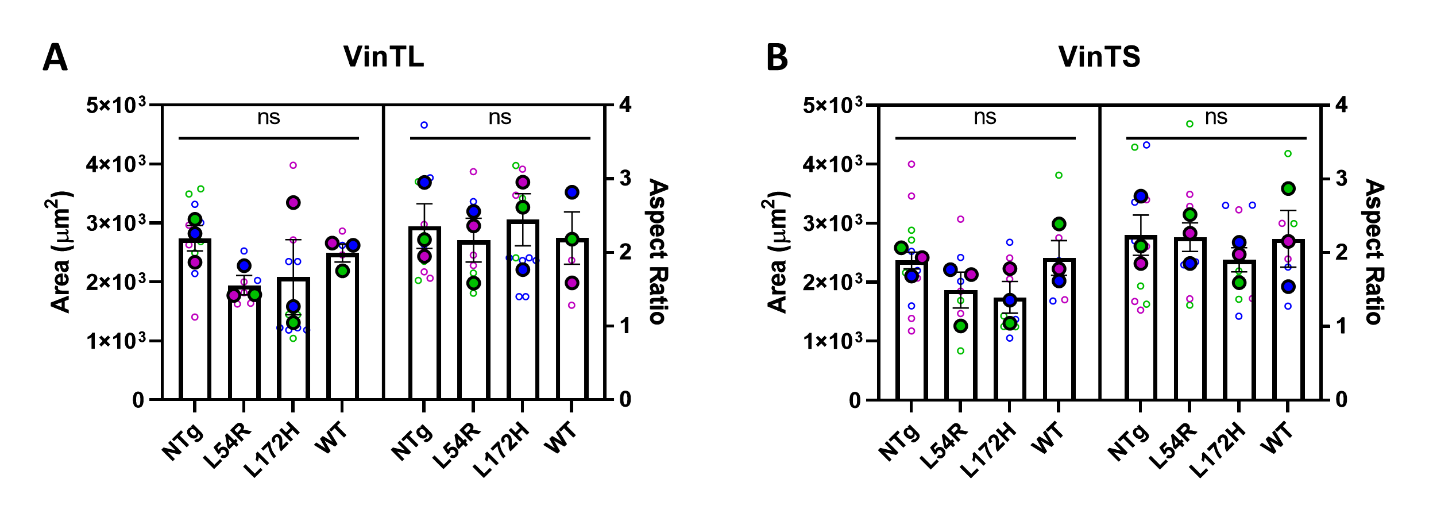
**Supplementary Figure 13. Cell morphology across cell lines.** Cell area and aspect ratio comparison for N=3 **A.** VinTL and **B.** VinTS transfected myoblast lines. Large dots denote the mean of independent experiments and smaller dots denote the mean of individual measurements. Each experiment and corresponding individual measurements are colored the same. Data analyzed via one-way ANOVA; ns, not significant. All error bars represent SEM.

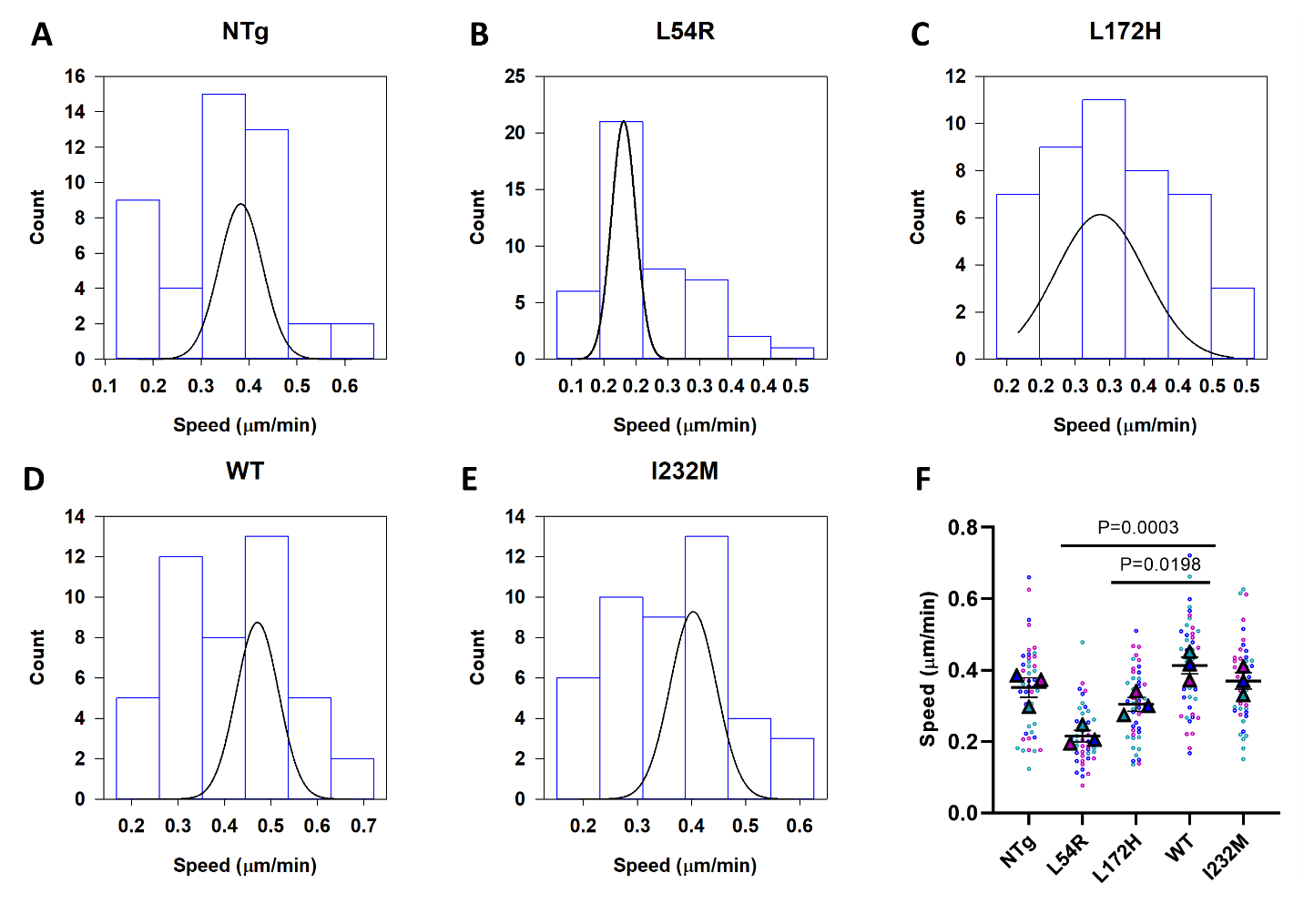

**Supplementary Figure 14. Single cell speeds.** Histogram of single cell speed distributions (N=3, 15 trajectories each) fitted to a non-linear normal regression for **A.** NTg, **B.** L54R, **C.** L172H, **D.** WT and **E.** I232M. **F.** Comparison of average speeds for each cell trajectory (small dots) and the mean speed of independent experiments (large dots). Data analyzed via one-way ANOVA; ns, not significant. All error bars represent SEM.

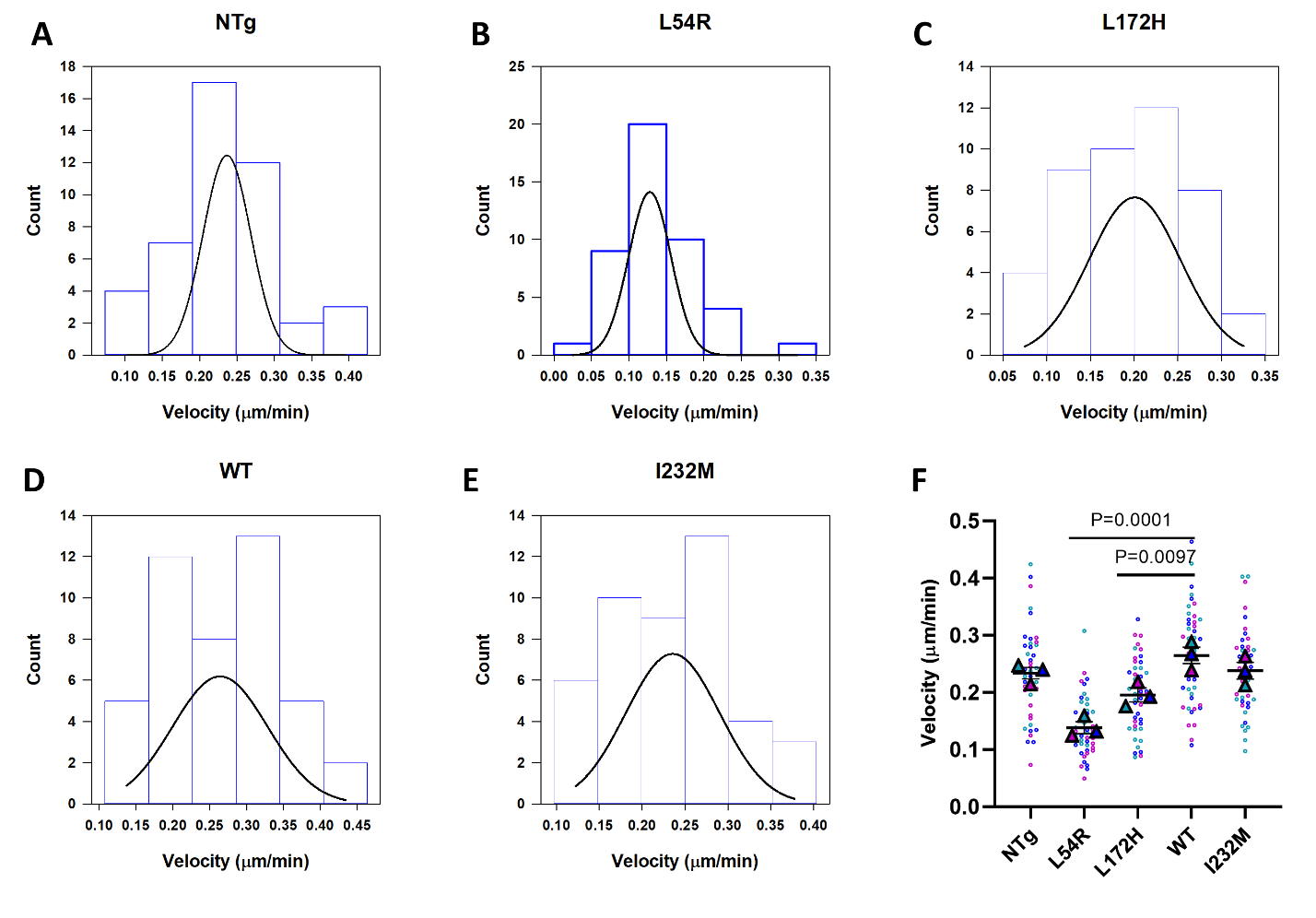

**Supplementary Figure 15. Single cell velocities.** Histogram of single cell velocities distributions (N=3, 15 trajectories each) fitted to a non-linear normal regression for **A.** NTg, **B.** L54R, **C.** L172H, **D.** WT and **E.** I232M. **F.** Comparison of average velocities for each cell trajectory (small dots) and the mean speed of independent experiments (large dots). Data analyzed via one-way ANOVA; ns, not significant. All error bars represent SEM.

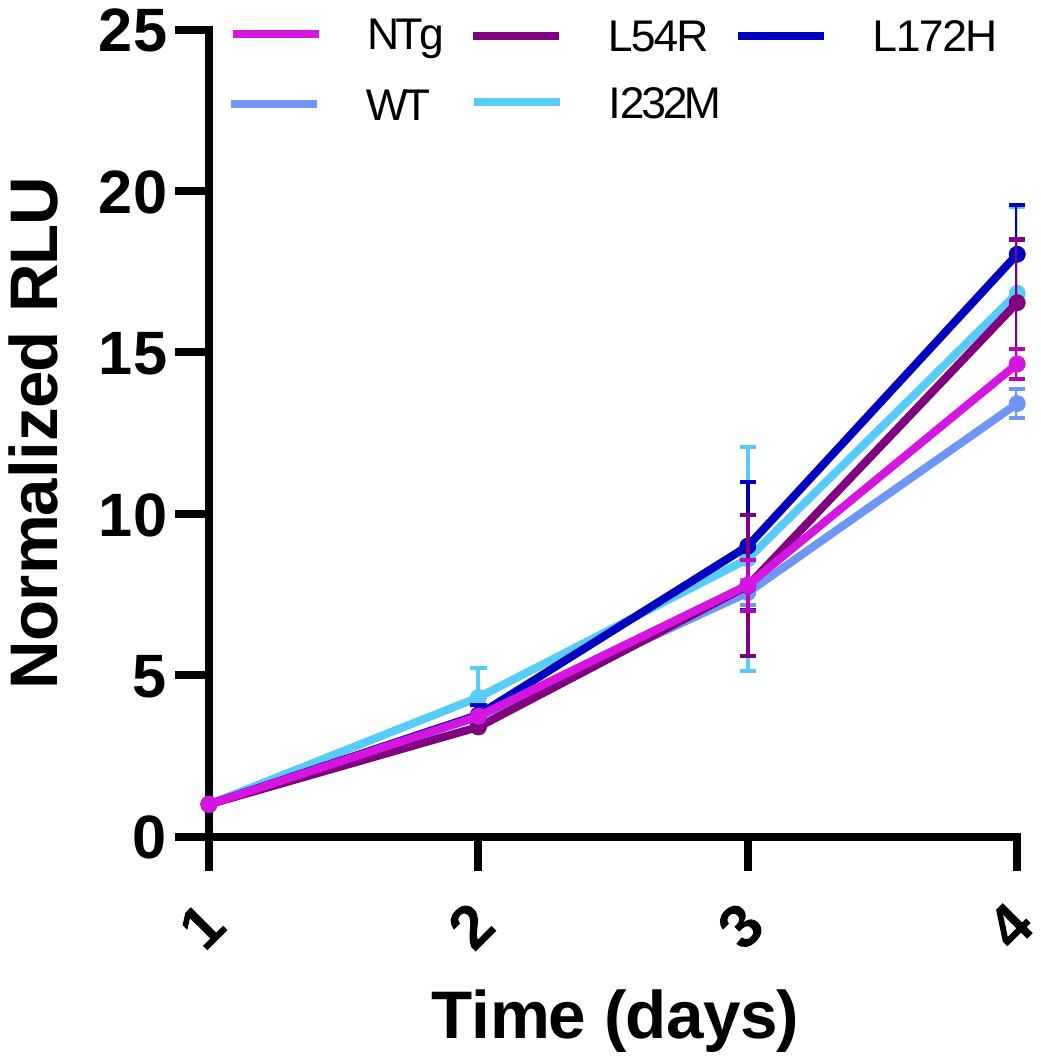

**Supplementary Figure 16. Cell proliferation curve.** CellTiter-Glo® luminescent cell viability assay to reflect cell proliferation (N=3), shown as normalized relative luminescent units over a period of 4 days.
